# A single-cell level and connectome-derived computational model of the *Drosophila* brain

**DOI:** 10.1101/391474

**Authors:** Yu-Chi Huang, Cheng-Te Wang, Ta-Shun Su, Kuo-Wei Kao, Yen-Jen Lin, Ann-Shyn Chiang, Chung-Chuan Lo

**Affiliations:** Institute of Systems Neuroscience, National Tsing Hua University, Hsinchu 30013, Taiwan; Brain Research Center, National Tsing Hua University, Hsinchu 30013, Taiwan; National Center for High-performance Computing, Hsinchu 30076, Taiwan; Department of Biomedical Science and Environmental Biology, Kaohsiung Medical University, Kaohsiung, Taiwan; Institute of Physics, Academia Sinica, Nankang, Taipei 11529, Taiwan; Institute of Molecular and Genomic Medicine, National Health Research Institutes, Zhunan, Miaoli 35053, Taiwan; Kavli Institute for Brain and Mind, University of California at San Diego, La Jolla, CA, USA

## Abstract

Computer simulations play an important role in testing hypotheses, integrating knowledge, and providing predictions of neural circuit functions. While considerable effort has been dedicated into simulating primate or rodent brains, the fruit fly *(Drosophila melanogaster)* is becoming a promising model animal in computational neuroscience for its small brain size, complex cognitive behavior, and abundancy of data available from genes to circuits. Moreover, several *Drosophila* connectome projects have generated a large number of neuronal images that account for a significant portion of the brain, making a systematic investigation of the whole brain circuit possible. Supported by FlyCircuit (http://www.flycircuit.tw), one of the largest *Drosophila* neuron image databases, we began a long-term project with the goal to construct a whole-brain spiking network model of the *Drosophila* brain. In this paper, we report the outcome of the first phase of the project. We developed the Flysim platform, which 1) identifies the polarity of each neuron arbor, 2) predicts connections between neurons, 3) translates morphology data from the database into physiology parameters for computational modeling, 4) reconstructs a brain-wide network model, which consists of 20,089 neurons and 1,044,020 synapses, and 5) performs computer simulations of the resting state. We compared the reconstructed brain network with a randomized brain network by shuffling the connections of each neuron. We found that the reconstructed brain can be easily stabilized by implementing synaptic short-term depression, while the randomized one exhibited seizure-like firing activity under the same treatment. Furthermore, the reconstructed *Drosophila* brain was structurally and dynamically more diverse than the randomized one and exhibited both Poisson-like and patterned firing activities. Despite being at its early stage of development, this single-cell level brain model allows us to study some of the fundamental properties of neural networks including network balance, critical behavior, long-term stability, and plasticity.

## Introduction

Understanding brain function requires knowledge of both molecular biology at the cellular level and of the interactions between neurons and the underlying circuit structure(Morgan and Lichtman 2013). In addition to various experimental approaches, computational modeling is becoming an increasingly important technique because it facilitates the validation of hypotheses and theories regarding neural circuit operation through the integration of existing observations into computer models(Churchland and Abbott 2016; Chaudhuri and Fiete 2016; Denève and Machens 2016; Sporns 2013). Indeed, extensive studies on neural network models covering *Caenorhabditis* elegans(Szigeti et al. 2014; Izquierdo and Beer 2016; Palyanov et al. 2011), insects(Wessnitzer and Webb 2006), rodents, and primates(Markram 2006; Izhikevich and Edelman 2008; Eliasmith et al. 2012) have greatly contributed to our understanding of neural circuit functions at the systems level. However, computer modeling also faces two major challenges: 1) a large number of neural network models were built to simulate specific functions in one or few brain regions(Izhikevich and Edelman 2008; Eliasmith et al. 2012). This approach limits our ability to study integrated functions or behavior at the systems level. 2) Due to the lack of connectomic data at the single-cell level for most species, large-scale neural network models can only be constructed based on the connectome at the macroscopic level(Izhikevich and Edelman 2008).

These challenges can be addressed by large-scale connectome projects(Milham 2012; Burns, Vogelstein, and Szalay 2014; Peng et al. 2015; Lo and Chiang 2016), which aim to reconstruct a high-resolution connectome of the whole brain at the single-cell level. While this is still a major challenge for large animals such as primates(Helmstaedter 2013), acquisition of single-cell level connectomes for small animals, such as the *Drosophila melanogaster* (fruit fly), has seen rapid progress(Chiang et al. 2011; Takemura et al. 2013; Shinomiya et al. 2011). Therefore, we suggest that the *Drosophila* is currently one of the best model animals for developing a high-resolution full brain computational model due to the availability of extensive neuron databases and neuroinformatics tools (Chiang et al. 2011; Shinomiya et al. 2011; Osumi-Sutherland et al. 2012; Parekh and Ascoli 2013; Ukani et al. 2016; Givon and Lazar 2016; Zheng et al. 2018; “Frontiers | Neuroarch: A Graph-Based Platform for Constructing and Querying Models of the Fruit Fly Brain Architecture” n.d.). Although being relatively small and simple, the fruit fly brain exhibits many high-level functions, including learning, memory, pattern recognition, decision making, and others. Hence, studying the neural circuits of small animals (insects) is extremely useful for our understanding of many essential brain functions(Webb and Wystrach 2016; Wessnitzer and Webb 2006; Su et al. 2017; Chang et al. 2017), and constructing a full brain model of the fruit fly brain may enable us to investigate how different subsystems in the brain integrate and how high-level behavior is carried out.

In this paper, we present our result from the first phase (Figure 1) of the Flysim project, a long-term research project aiming to develop a full-brain computational model of the *Drosophila* brain at the cellular and synaptic levels. The most distinct difference between the proposed model and other large-scale brain models is that in the proposed model every neuron was uniquely derived from a neuron image from the FlyCircuit database(Chiang et al. 2011). The database currently hosts 28573 and 22835 neuron images from female and male *Drosophila* brains, respectively, and the amount of data is rapidly increasing. The 22835 images account for 22.83%-15.22% of the estimated total neurons (100,000-150,000) in a *Drosophila* brain. Although being a small percentage, these neurons fairly represent the entire brain as they widely distributed throughout every neuropil and cover more than 93% voxels (each voxel is 0.32×0.32×0.64 μm in dimension) of the standard brain space (Chiang et al. 2011).

**Figure 1.**
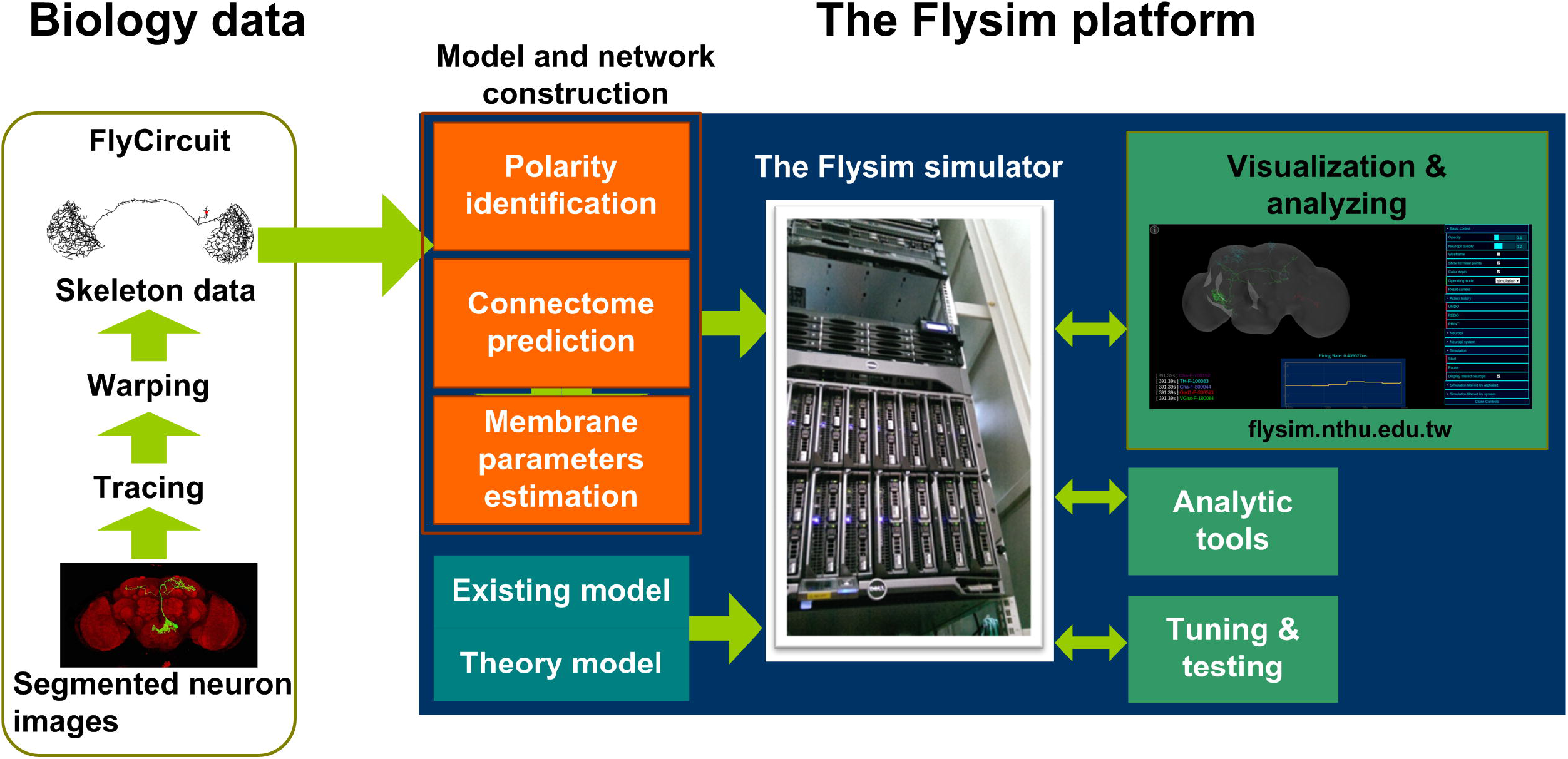
The Flysim platform for *Drosophila* full-brain modeling. The platform imports neuron skeleton data from the FlyCircuit database. The data undergo several processes before they are transformed into a computational model. The processes include polarity (axon or dendrite) identification, neuron connection prediction, and membrane parameter estimation. These processes lead to a raw model that can be simulated by the Flysim simulator, developed in-house. The raw model requires tuning and testing before it reaches a stable state. The activity of the stabilized brain model can be visualized on the web and used for studying brain dynamics.

Reconstructing a full-brain model based on a neuron image database poses several challenges. In the first phase of the project we developed mathematical and statistical tools that are required for transforming the neuronal morphologies into computational models and for deriving parameters that allow the modeled brain to maintain a stable resting state. Specifically, we needed to 1) predict the polarity of each neuron based on its morphology, 2) infer the synaptic connections and their weight between any two neurons, 3) derive membrane parameters for each neuron based on its size, 3) design a neural network simulator that is able to accommodate the simulations, and 4) find the balance condition of the brain model that is active and stable in the resting state. We also analyzed the network structure and the activity of the reconstructed fruit fly brain and found that it exhibits much higher diversity yet more stability than those observed in a randomized brain network. Finally, we discuss the issues in the current model, including identification of neuron type, receptor type and polarity, models for modulatory synapses, image alignment, and choice of single neuron model. We further suggest the technology and methodology that are required to address these issues in the next phase of the model development.

## Material and method

### Data preprocessing and analysis

The FlyCircuit database provides detailed neuron images and accurate tracing lines (skeletons) for each neuron. However, to construct a computational model of the brain network, we need the following additional information: 1) polarity of each neuron arbor, 2) connections between neurons, and 3) their physiological properties. Here we describe the methods we used to estimate the parameters associated with these properties.

#### Synapse polarity prediction and validation

The information regarding the polarity (axon and dendrite) of each neuron was not available in the original neuron skeleton data obtained from the FlyCircuit database. To infer the polarity, we used the SPIN method(Lee et al., n.d.), which is a machine-learning algorithm designed for identifying the axonal and dendritic domains of a neuron based on its skeleton. Although this method is not 100% accurate (~84-92% on the original test dataset (Lee et al., n.d.)), it is the only available automated method that can be applied to a large-scale neuron image database.

The original SPIN method was tested on a small subset of neurons that innervate the protocerebral bridge (PB) and modulus (MD). To apply this method to the entire brain, we tweaked several parameters and re-trained the classifier. We first randomly selected 90 neurons that cover diverse morphologies from several neuropils including the PB, MD, antennal lobe (AL), and mushroom body (MB). We chose these neuropils because the polarity of their neurons is largely known. We manually labeled the polarity of the neurons and used them as the training data for SPIN. To identify the best combination of the morphological features for polarity classification, we tested all three feature selection methods provided by SPIN: sequential, exhaustive, and manual assignment. We found that the sequential method provided the best result, which indicated that there are 11 morphological features correlated with the polarity (Table 1). Among the 11 features, the top five are: path length to soma, mean branch order, maximum path length, maximum branch order, and number of branch points and volume of the convex hull.

**Table 1.**
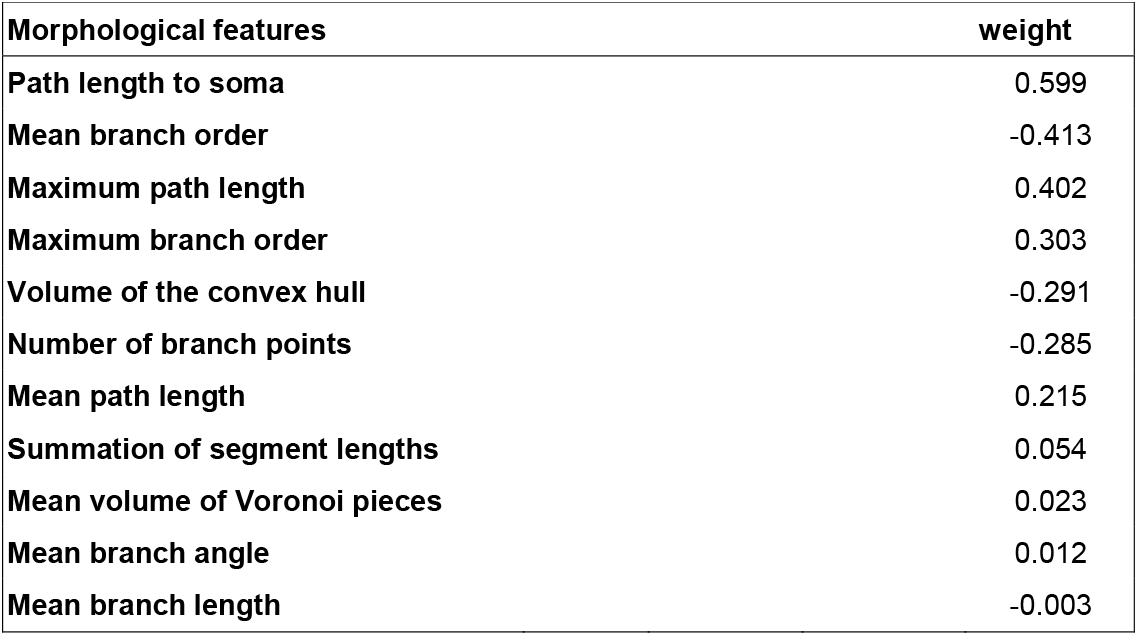
Morphological features that are correlated with the neuronal polarity as determined by the SPIN method. The weight represents the degree of correlation. Positive values indicate positive correlation while negative values indicate negative correlation. The definition of each feature is described in Cuntz et al. 2010(Cuntz et al. 2010).

| Morphological features | weight |
| --- | --- |
| Path length to soma | 0.599 |
| Mean branch order | -0.413 |
| Maximum path length | 0.402 |
| Maximum branch order | 0.303 |
| Volume of the convex hull | -0.291 |
| Number of branch points | -0.285 |
| Mean path length | 0.215 |
| Summation of segment lengths | 0.054 |
| Mean volume of Voronoi pieces | 0.023 |
| Mean branch angle | 0.012 |
| Mean branch length | -0.003 |

The training yielded a new polarity classifier. Next, SPIN separated each test neuron into several domains and classified the polarity of each domain. Because the data used in the present study have a higher resolution, i.e., more terminal points, than those used in the development of the SPIN method, SPIN tended to separate some neurons into too many domains. This issue was resolved by changing the parameter Th_DP_ from 0.01 to 0.001. To validate the performance of the new classifier, we selected the 442 neurons that were reported in Lin et al 2013(Lin et al. 2013) as test neurons because their polarity has been reported in detail by two studies(Lin et al. 2013; Wolff, Iyer, and Rubin 2015). We removed the EIP neuron class, which innervates the ellipsoid body, inferior dorsofrontal protocerebrum, and protocerebral bridge, because the reported polarity is inconsistent between the two studies. Our test result indicated a 91.3% of terminal level accuracy, which means that on average, the polarity of 91.3% of the terminals in each neuron was correctly classified. Finally, we used the new classifier to classify the polarity of all the neurons in the FlyCircuit database.

#### Synapse weight prediction and connection validation

Next, we estimated whether connections exist between any two given neurons. In FlyCircuit, each neuron image was taken from one individual fly brain. Although each image has been transformed (or warped) and registered in a standard brain space, this process inevitably created warping error. Ideally, the probability of synapse formation between two neurons is correlated with the degree of contact between them (Douglass and Strausfeld 2003; Tanaka, Endo, and Ito 2012; Peters and Payne 1993). However, due to the warping error, if two neurons have closely contacted branches in the standard brain space, this does not necessarily indicate that they form synapses. Likewise, two neurons that are not closely contacted in the standard brain space may in fact form synapses (Figure 2A and 2B). Therefore, additional procedures were required in order to infer the probability of synaptic formation between neurons.

**Figure 2.**
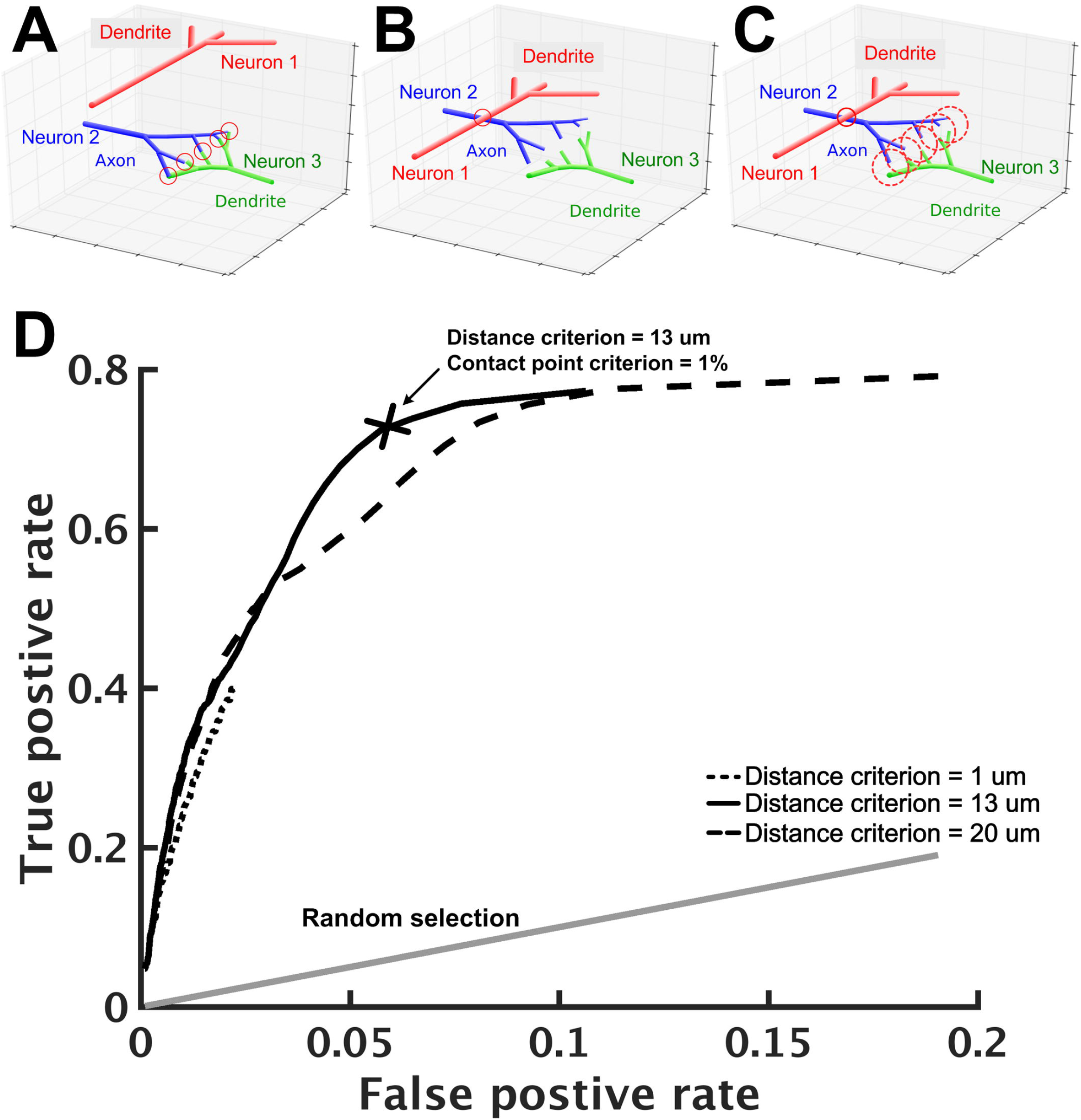
Prediction of neuron connections based on distance and number of contact points. (A) - (C) Schematics of neuron connections illustrate how the prediction error can be reduced by the consideration of distance and contact points. (A) The dendritic arbor of neuron 1 is far apart from the axonal arbor of neuron 2 and they do not form any synapse. Neuron 2 and neuron 3, however, form five synapses as indicated by the five contact points (red circles) between them. (B) Warping error may occur when neurons are transformed and aligned in the standard brain space. In this case, neurons 1 and 2 come in contact while neurons 2 and 3 become separated. If the connection prediction is made only based on the distance between neuron processes, errors would occur in this case. (C) To address this issue, we set two criteria: contact point and distance. Axonal and dendritic branches are counted as having a contact point when their shortest distance falls within a preset distance. Two neurons are considered to form synapses when their contact point number is larger than a preset value. When proper values for the two criteria are set, neurons 1 and 2 are no longer connected but neurons 2 and 3 become connected. (D) Using the receiver operating characteristic analysis with various contact point and distance criteria, we identified the best criteria that lead to a high true positive rate (x-axis) with a reasonably low false positive rate (y-axis). Each black line represents a fixed distance criterion (dot: 1 *μ*m, solid: 13 *μ*m, dashed: 20 *μ*m) with varying contact point criteria. The gray line represents the result when synaptic connections between neurons are randomly assigned. The cross indicates the best criteria: the contact point number > 0.1% (of the total input contact points of the presynaptic neuron, see text) and the distance < 13 *μ*m.

To this end, we designed a protocol that infers neuronal connections based on two criteria: distance and contact point. The distance criterion sets a maximum distance between an axonal segment of one neuron and a dendritic segment of another neuron that can be considered to be forming a contact point. A segment is the straight line between two consecutive nodes on a neuronal skeleton. For two selected neurons, we calculated the distances for all pairs of segments (one from each neuron) with different polarity. Next, we counted the number of contact points for these two neurons. The contact point criterion sets the minimum number of contact points between two neurons that can be considered to be forming synapses (Figure 2C). We used the relative number *R*, rather than the absolute number, for the contact point criterion. Specifically, if *N_ik_* represents the number of contact points between neuron *i* (axonal side) and neuron *k* (dendritic side), then neuron *i* is considered as forming synapses with neuron *k* if 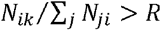. Intuitively, one would place the number of all output contact points, i.e., Σ*_j_ N_ij_*, in the denominator, so that *R* represents the ratio between the contact points of neuron *i* to neuron *k* and the contact points of neuron *i* to all downstream neurons. However, such a ratio leads to an undesired consequence, which limits the possible number of downstream neurons. For example, if *R* is set to 0.01, neuron *i* will have no more than 99 downstream neurons. This is because if we rank the downstream neurons by their contact points with neuron *i*, the 100th downstream neuron must have an *R* smaller than 0.01. This problem will have a strong impact on neurons that have a large number of downstream neurons. Instead, using the number of all input contact points, i.e., Σ*_j_ N_ji_* as the denominator solves the problem. Although it seems odd to calculate the ratio based on the number of input contact points, it is not because the number is in fact roughly proportional to its total number of output contact points (S1 Fig).

The optimal values of *D* and *R* for the two criteria were determined by the following procedure: 1) we started from a small distance criterion (*D* = 1 μm) and contact point criterion (*R* = 0.1%), 2) for every pair of neurons in the test neuron set, we calculated the number of contact points and determined whether the two neurons form synapses based on the criteria; 3) we compared the result with data from a previous research (Lin et al. 2013) and calculated the true positive rate and false positive rate, and 4) we changed the distance and the contact point criteria and repeated steps 2–3. Finally, we used the receiver operating characteristic (ROC) analysis (Fawcett 2006; Lasko et al. 2005) to determine the best criteria to be: distance = 13 *μ*m with contact points = 1% (Figure 2D). With these criteria, we achieved an acceptable true positive rate of 0.71 and a very low false positive rate of 0.058.

All procedures were performed with the 442 neurons reported in Lin et al 2013(Lin et al. 2013). Based on Lin et al 2013(Lin et al. 2013) and Wolff et al. 2015(Wolff, Iyer, and Rubin 2015), who reported the anatomy of the same circuits, we were able to derive the network connections of these neurons and use them as a reference to optimize our connection estimation protocol. Lin et al 2013(Lin et al. 2013) and Wolff et al. 2015(Wolff, Iyer, and Rubin 2015) reported the polarity and innervated subregions of each neuron. To construct the reference connectivity of these neurons, we assumed that a neuron that projects its axonal arbor to a glomerulus forms synapses with another neuron that has its dendritic arbor in the same glomerulus. Our assumption is reasonable considering that each defined glomerulus takes a small spatial volume (on average 16 μm in size(Chang et al. 2017)) and a neuron that innervates a subregion typically fills up the volume with its arbors and highly overlaps with other innervated neurons.

#### Estimate of Membrane parameters

For each neuron, we estimated its membrane parameters in order to create a LIF model for simulation. The LIF model requires the following parameters: resting potential *V*_resting_, spike threshold *V*_threshold_, reset potential *V*_reset_, refractory period *T*_refract_, membrane time constant τ_m_, and membrane capacitance *C*_m_. To determine the first three parameters, we extensively reviewed the literature and estimate the typical value for each parameter (S1 Table). In consequence, we set *V*_resting_ = -70 mV, *V*_threshold_= -45 mV, *V*_reset_ = -55 mV, and τ_m_ = 16 ms for every neuron. The refractory period was set to 2.0 ms. The membrane capacitance of each neuron was size-dependent and was determined by the following procedures.

The membrane capacitance, *C*_m_, depends on the total area of the cell and hence roughly correlates with the size, or the total branch length, of the cell. Therefore, at the current stage we simply assumed that *C_m_* of a cell linearly correlates with its total skeleton length. Based on this assumption, we can easily estimate the *C*_m_ for each neuron if we find the typical value of the membrane capacitance per unit length of the skeleton, denoted *c*_m_. Although this was a very rough estimate, it gave us a size-dependent membrane capacitance and is certainly superior to simply setting all neurons with an equal membrane capacitance. We have found that cell membrane capacitance was 0.6 μF/*cm*^2^ to 1.0 μF/*cm*^2^ from previous work (Gouwens and Wilson 2009)(Weir, Schnell, and Dickinson 2014) and we considered the average value, 0.8μF/*cm*^2^, as the membrane capacitance per unit area for our neuron model. Because the information about the diameter of each neuron branch is not available in the current database, we were not able to directly calculate the membrane area of a neuron but had to estimate the value indirectly based on other studies. Wilson and Laurent 2005(Wilson and Laurent 2005) measured the total length and area of three antennal lobe projection neurons. By comparing the skeleton length of the neurons in our database to that reported by Wilson and Laurent 2005(Wilson and Laurent 2005), we obtained an empirical equation for the total area *A* of a neuron, *A* = (*l_i_*×2*π*× 0.147) × 2.38 + 5340, where *l_i_* is the skeleton length of the neuron *i*. By multiplying *A* by 0.8 μF/*cm*^2^, we obtained the estimated membrane conductance of each neuron.

### Model network construction

Based on the procedures describe above, we established a brain-wide neural circuit model including an individual LIF model (described below) for each neuron and the conductance-based synapses formed by these neurons. We acquired neurons from the female fruit flies in the FlyCircuit database, and excluded the isolated neurons (those not connected to any other neurons based on our connection estimation). We obtained a total of the 20,089 neurons that can be used in the brain-wide circuit model. Next, we inferred the type, in terms of released transmitters, of each neuron by the driver used to image the given neuron. The driver type is indicated by the first part of a neuron’s name in the database. For example, the neuron named VGlut-F-200532 is assumed to be a glutamatergic neuron. Specifically, there were 3365 putative cholinergic (Cha) neurons, 5998 putative glutamatergic (VGlut) neurons, and 7956 putative GABAergic (Gad) neurons. At the current stage we only simulated synaptic projections from these three types of neurons, which form a total of 1,044,020 synapses. The other 2,770 neurons were likely modulatory neurons, which release neurotransmitters such as dopamine, octopamine, serotonin, and others. We argue that it is safe to exclude their synapses at the current stage because their slow effect does not significantly impact brain dynamics at the millisecond to second time scales, as the present study focused on. We will include the modulatory synapses in the future when we simulate the fruit fly behavior at the minute to hour time scales.

#### Neuron and synapse models

Each neuron was simulated by a compartment of the LIF model with conductance-based synapses. The neuron model is described by:

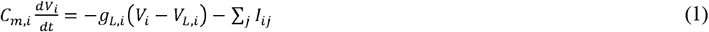

where the subscripts *i* and *j* are the neuron indices, g_L_ = *C_m_/τ_m_* is the membrane conductance, *V_L_* (= *V*_resting_) is the resting potential, and *I_ij_* is the synaptic current, which is contributed by glutamatergic (including AMPA and NMDA receptors), cholinergic (Ach), and GABAergic (GABA_A_) synapses formed by projections from the presynaptic neuron *j*. For AMPA receptors in glutamatergic synapses, as well as cholinergic and GABAergic synapses, we have

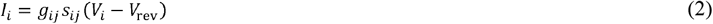

and for NMDA receptors in glutamatergic synapses, we have

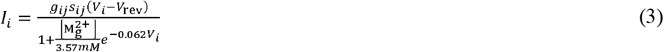

where *g* and *s* are the synaptic conductance and the gating variable, respectively, *V*_rev_ is the reversal potential, which is 0 mV for the excitatory (including AMPA, NMDA, and Ach) and -70 mV for the inhibitory (GABA_A_) synapses, respectively, and [Mg^2+^] (=1mM) is the extracellular magnesium concentration, which is used to describe the effect of the magnesium block on the NMDA channels. We would like to clarify the use of the term “synapse.” In biology, a neuron can make multiple contacts and form multiple synapses with another neuron. However, in the single-compartmental model used in the present study, the effect of multiple synapses between two neurons can be combined and described by only one synaptic equation. Therefore, a model synapse between the presynaptic neuron *i* and the postsynaptic neuron *j* can be treated as a collection of all biological synapses formed between the two neurons, *i* and *j*. The gating variable *s_ij_* is given by:

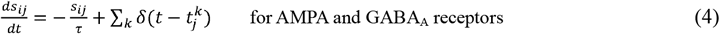

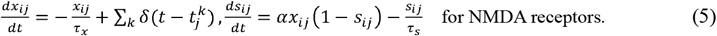

*δ* is the delta function, and 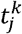 is the time of the *k*-th spike from the presynaptic neuron *j*. The synaptic conductance *g* is an unknown parameter that indicates the strength of the synapse. We assume that the synaptic strength between a presynaptic neuron *i* and postsynaptic neuron *j* are proportional to the number of their contact points, *N_ij_*

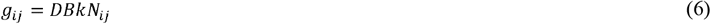

The proportion constant is the multiplication of three variables: *D*, *B*, and k. *D* is a variable for short-term depression described below. *B* was different between excitatory and inhibitory synapses and was used to adjust the balance between excitation and inhibition of the system as described in the Results. *k* is a variable used to balance the relative contribution between excitatory synapses that contain AMPA, NMDA, or Ach receptors. *k* was set to be 1/300 for AMPA receptors. Because the NMDA time constant is 50 times larger than that of AMPA, we set *k* = 1/15000 for NMDA receptors, so that both NMDA and AMPA contributed equally to the synaptic current in a glutamatergic synapse. Likewise, *k* was set to be 1/3000 for an Ach synapse because its time constant is 10 times larger than that of AMPA. For GABA_A_ synapses, *k* was set to be 1/300, which is equal to that of AMPA. Note that for a given glutamatergic synapse, the corresponding NMDA and AMPA components shared the same *D*, *N_ij_*, and *B*.

We delivered to each neuron a small but fluctuating membrane current as the background noise. Specifically, at each time step and for each neuron, a value of membrane current was drawn from a Gaussian distribution and was applied to the neuron in order to generate membrane potential fluctuation. The width of the Gaussian distribution is dependent on the size of each neuron to ensure that the resulting mean membrane potential (= -60 mV) and its standard deviation (3 mV) at the resting state are the same for all neurons. The background noise is so small that each neuron barely fires (with a mean firing rate of ~0.004 Hz) without external synaptic input.

#### Short-term plasticity

We implemented the STD, a feature commonly observed in various nervous systems including the *Drosophila*’s (Wilson and Laurent 2005; Nagel, Hong, and Wilson 2015; Root et al. 2007). We adopted a model which describes STD as a presynaptic calcium dependent dynamic, in which the available vesicles decrease following each presynaptic spike and exponentially return to the baseline with a long time constant(Hempel et al. 2000; Varela et al. 1997; Abbott et al. 1997). Specifically, the STD variable *D* is given by:

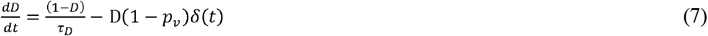

where τ_D_ is the time constant of STD, and *p_v_* is the synapse vesicle release probability(X.-J. Wang 1999), δ(t) is a delta function that is infinity at the time of every presynaptic spike and 0 elsewhere. *D* is used to modulate the synaptic conductance as indicated in Eq. (6).

### The randomized brain network

To investigate the neural network dynamics of the reconstructed fruit fly brain, it is useful to compare it to a randomized network to assess the contribution of the brain network structure to the network dynamics. To this end, we created a randomized fruit fly brain network using the following procedure. We preserved all neurons in the reconstructed fruit fly brain model as well as all synaptic conductance *g_ij_*’s. Next, we rewired all connections by randomly assigning a new postsynaptic neuron *i* to every *g_ij_*, while keeping the presynaptic neuron *j* unchanged. The randomized fruit fly brain network had the same number of neurons, the same number of synapses, and the same synaptic weight (*g_ij_*) distribution with those in the reconstructed fruit fly brain network. Because of the random rewiring, the isolated neurons in the reconstructed brain network became connected in the randomized brain network, which had a slightly larger number of neurons (22,835).

### Model network simulation

To perform simulations for the model fruit fly brain, we built a neural network simulator, Flysim, in C++. Flysim includes four major components: 1) two-pass network compilation, 2) data managing and optimization, 3) computation, and 4) data output.

#### Two-pass network compilation

The network building process requires a special design because of the large size of the parameter file, which specifies unique parameters for each of the 20,089 neurons and 1,044,020 synapses. In order to facilitate the computer memory access and to shorten the network construction time in this large-scale neuron network, we utilized the “two-pass compiler” concept in network compilation. In the first pass (Figure 3A), Flysim reads through the parameter file, calculates the number of total neurons and synapses, and allocates the memory for each neuron and synapse. In the second pass (Figure 3B), Flysim reads every parameter and fill them into the pre-allocated memory. This two-pass approach avoids the time needed for dynamic memory allocation when building neuron data, and hence reduces the time for network construction from over 15 minutes down to only 1.5 minutes.

**Figure 3.**
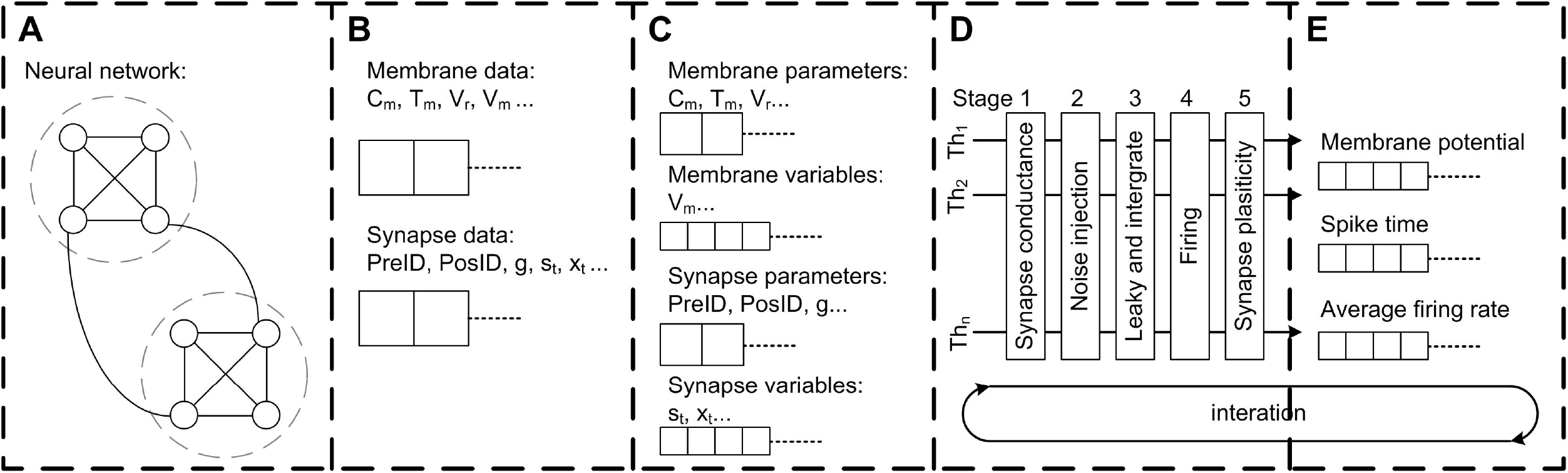
The architecture of the Flysim simulator. In the fruit fly brain model, each neuron and synaptic connection are unique. Therefore, the entire model requires a large amount of computer memory. The simulator is designed to address this challenge. (A) The simulator first goes through the network configuration file and estimates the number of neurons and synapses. Next, the simulator pre-allocates memory. (B) The simulator goes through the network configuration file again, reads all parameters, and then builds the whole network by filling each membrane-related and synapse-related datum into the pre-allocated memory. (C) The simulator performs linear reduction for synapse-related data to reduce computation and archive threading level parallelism. (D) The simulator dispatches each thread with one assembled neuron array and aligns each thread into a 5-stage pipeline for parallel processing. (E) Simulation results, including spike, membrane potential, average firing rate, and other data are directly accessed from assembled neuron arrays to archive high throughput and low latency data output.

#### Data managing and optimization

We also adopted the compact data structure to reduce memory access. Because in our network model each neuron has different parameters and connections, the data are not linearly reducible. To improve the memory access efficiently, we separated the data into two categories: membrane-related and synapse-related (Figure 3C). Flysim sorts the synapse-related data, which are compiled in the previous process, and then reduces the fast-responding gating variables of each neuron as follows. In our network model, the dynamics of fast-responding receptors such as AMPA, GABA_A_, and acetylcholine are described by a simple exponential decay. This property makes it possible for us to linearly combine all gating variables of the same receptor type (AMPA, GABAa, or acetylcholine) in each post-synaptic neuron *i* into one single variable, *S_i_*:

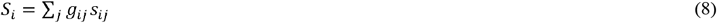

where *g_ij_* and *s_ij_* are defined in equations (4) and (6), respectively. The dynamics of are described by

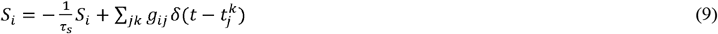

where 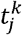 indicates the time of *k*-th spikes from the presynaptic neuron *j*. Instead of calculating a large number of gating variables for each connection for a given neuron, we only needed to calculate one gating variable for each receptor type. In the simulator, Equation 9 was used to replace Equation 4 for the AMPA, GABA_A_, and cholinergic receptors. This reduction led to program space and time localities, which greatly improved memory fetch and storage through the high-speed buffering mechanism in the modern computer memory hierarchy.

#### Computation

To further reduce the computation time, we entered the calculations of membrane current and potential in the same program block for spatial and temporal localities, which allowed the C++ compiler to automatically optimize the operations and improve the speed.

When performing threading level parallelism (TLP), load balance greatly influences computing performance (Figure 3D). Load balance can be easily achieved for neuron-related data because each neuron is described by the same number of neuronal parameters. However, this is not the case for synapse-related data because the number of synapses varies greatly between neurons. To address this issue, we assembled multiple arrays, and each contained synapse-related data from randomly selected neurons. Due to the nature of random selection, the arrays were roughly of the same size, or balanced. Each array was then loaded into one thread for computation. By performing TLP with load balance, we could achieve a 1:35 simulation speed (1 second of biological time requires 35 s of real time to simulate) using four threads with the current network size (20,089 neurons and 1,044,020 synapses) (see Results).

We found that the synaptic strengths in the reconstructed brain network have broad distributions. Therefore, some neurons received an extremely large number of innervations from GABAergic neurons, which produced excessive inhibitory current input and brought the membrane potential of the postsynaptic neuron to a level much lower than the reversal potential, or *V_i_* ≪ *V_rev_*, in equation 2. When this occurred, subsequent GABAergic input instead produced depolarized current (*I_i_* < 0, see equation 2). If the subsequent GABAergic input is again very strong, the large depolarization current might in fact bring the membrane potential above the firing threshold and generate an action potential. To eliminate such artifacts, we implemented a constrain on the maximum potential change *dV_max_* of a neuron in one simulation time step. The maximum value was set to be:

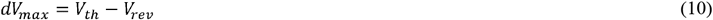

For the numerical solver, we used the first order exponential integrator method(Cox and Matthews 2002) instead of the commonly used 4th-order Runge-Kuta method. The reason is that due to the nature of the LIF model, we only needed to solve the equations for the sub-threshold membrane dynamics, which evolve much slower than those of spike activity. Using the exponential integrator can greatly improve the speed while at the same time retain high precision (comparable to the look-up table method). To speed up the generation of the Gaussian noise, which is used for membrane noise, we used the ziggurat(“The Ziggurat Method for Generating Random Variables | Marsaglia | Journal of Statistical Software” n.d.) method to generate a Gaussian-distributed random number. This approach further improved the speed three-fold compared to the standard C++ random number generator.

#### Data output

For the simulated data output module, we adopted a direct-access approach in which neuron variables are written to files directly rather than through a commonly used independent message queue or message buffer (Figure 3E). Flysim uses clock-driven simulation and it exports various data, including spike time, firing rate, membrane potential, and others, in each time step. The direct data-access approach provides a high output rate with low latency, and therefore minimizes the time spent on non-simulation processes.

## Results

### Statistics of the network structure

We first examined several key statistics of the reconstructed fruit fly brain network and found that it is highly diverse and exhibits interesting patterns of local connectivity. The network contains 20,089 neurons and the average number of edges (connections) is 52. The neuron sizes, as represented by individual neuron’s total skeleton length, cover two orders of magnitude. The distribution of neuron size forms two peaks, suggesting two distinct neuron types in the fruit fly brain (Figure 4A). Further analysis revealed that one peak mainly corresponds to the projection neurons (mean skeleton length = 1753 μm) and the other corresponds to the local neurons. Projection neurons are those innervating more than one neuropils and are usually much larger than the local neurons, which only innervate one neuropil. We further found that the local neuron distribution also formed two peaks. The peak that corresponds to the shorter mean length is mainly contributed by the local neurons in the medulla (MED), while the longer one is contributed by the rest of the local neurons (Figure 4A inset). The MED local neurons have a mean skeleton length of 858 μm, while the non-MED local neurons have a longer mean skeleton length of 1206 μm, which is still significantly shorter than that of the projection neurons (t-test, p < 10^-21^). We noted that the MED local neurons account for a significant number (1455) of the total neurons in our sample. However, considering that each MED consists of roughly eight hundred visual columns and each column contains a few dozen local neurons (Zhu 2013; Morante and Desplan 2008), the number of MED local neurons in our sample seems to be reasonable.

**Figure 4.**
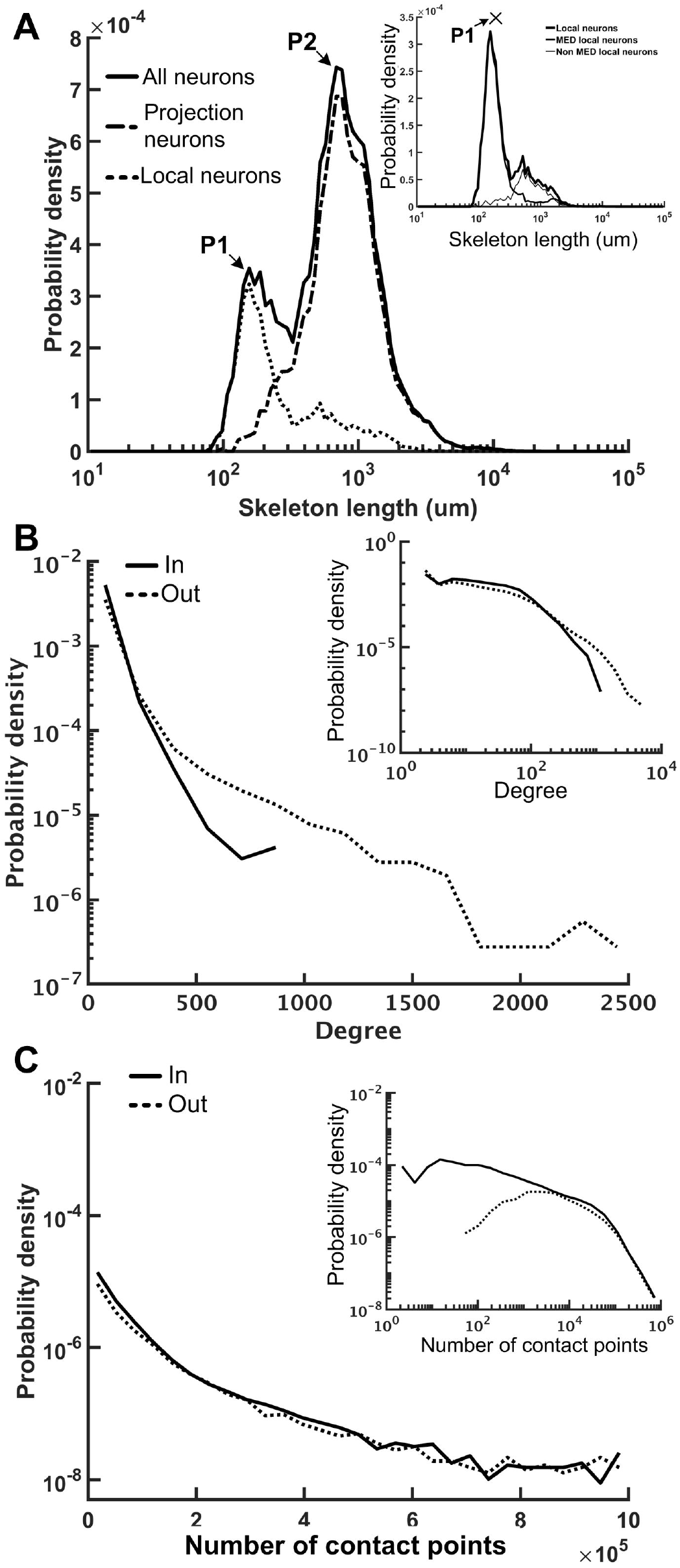
The neuron size and network connectivity of the fruit fly brain network are highly diverse. (A) the distribution of neuron size as represented by the total skeleton length. The probability density was calculated by dividing the number of neurons in each bin by the total number of neurons and by the bin size. The sizes for all neurons (thick black) exhibit a bi-modal distribution. The left peak is mainly contributed by the local neurons (dotted curve), while the right peak is mainly contributed by the projection neurons (dashed curve). The distribution of the local neurons also forms two peaks with the shorter-length peak contributed by medulla (MED) local neurons and the longer-length peak, by non-MED local neurons (inset). (B) The distribution of degree (number of connections of each neuron) follows a broad distribution for both in-degree (input connections) and out-degree (output connections). Inset: a double-log plot of the same curves. (C) The distribution of the contact point number of each neuron also exhibits a long tail distribution for both input and output contact points. Inset: the same curves in a double-log plot.

We further examined the connectivity of the fruit fly brain networks. The connectivity exhibited long tail distribution and connectivity was 0.003, meaning that each neuron made connections to ~0.3% of neurons in the brain, on average. The degree distribution, or the distribution of the number of connections made by each neuron, formed a broad distribution with the largest connection number up to 944 for in degree (input connections) and 3,982 for out degree (output connections) (Figure 4B). Both distributions roughly followed an exponential form, at large degrees. If we consider the full brain (estimated 100,000-150,000 neurons in total), connectivity of 0.3% gives rise to an average degree of 390 per neuron. Although the number seems to be high, note that the degree distribution follows a long distribution with a fat tail, suggesting that the average number is strongly influenced by a small number of highly connected neurons while the degrees of most neurons are smaller than 390.

Next, we examined the total input and output contact points of each neuron (see Methods) and found that their distribution also formed broad distributions, but with power-law tails (Figure 4C). The broad degree and connection weight distributions indicate that the connectivity of the fruit fly brain network is multi-scaled.

We further investigated the local connectivity under the consideration of neuron types, which influence the network balance. The fruit fly brain network, just like any other neural network, is characterized by strong recurrent/feedback connections with both excitatory and inhibitory synapses. We expect that the ratio between the excitatory and inhibitory input has to remain balanced. Otherwise, slightly more excitation (or less inhibition) could be quickly magnified through the recurrent connections and destabilize the entire network. A balanced network does not imply that it is lack of spontaneous activity or is unresponsive to the input as one may imagine. Several theoretical studies suggested that a balanced state can improve functionality of a neural network compared to unbalanced one (Lo, Wang, and Wang 2015; C.-T. Wang et al. 2013; Vogels and Abbott 2009; Chance, Abbott, and Reyes 2002) and such a balanced state has been observed in various nervous systems (Haider et al. 2006; Mariño et al. 2005; Shu, Hasenstaub, and McCormick 2003; Berg, Alaburda, and Hounsgaard 2007). We calculated the E-I index for each neuron and plotted its distribution separately for each neuron type (Figure 5). The E-I index of a neuron is defined as (*N_E_* − *N_I_*)/(*N_E_* + *N_I_*), where *N_E_* is the total excitatory input (from VGlu and Cha neurons) and *N_I_* is the total inhibitory input (from Gad neurons) to the given neuron. The E-I index can be calculated with unweighted or weighted input: the former only counts the number of input neurons and the latter weights each input with its contact point number.

**Figure 5.**
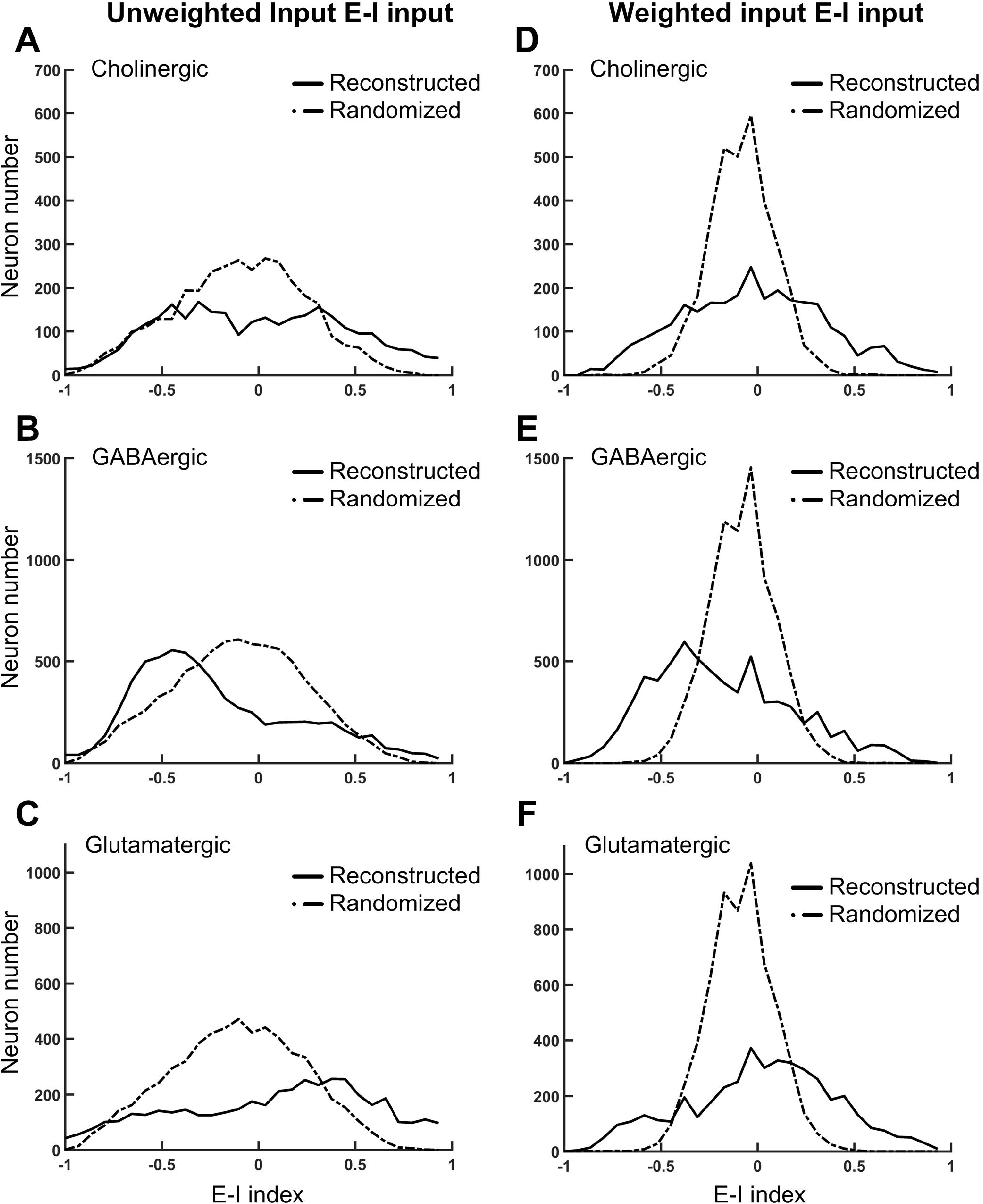
The normalized difference between the excitatory and inhibitory inputs (the E-I index) for different types of neurons in the fruit fly brain indicates the striking diversity of the local circuits. (A) - (C) The distributions of the E-I index of each neuron, plotted separately for the putative cholinergic, GABAergic, and glutamatergic neurons, respectively. (D) - (F). Same with the panels (A) - (C), respectively, but the E-I index are calculated based on the connections weighted by the contact point numbers. Solid curves: the reconstructed fruit fly brain. Dashed curves: the randomized fruit fly brain. The reconstructed brain exhibits much broader distributions than the randomized brain does in all conditions. The putative GABAergic neurons receive more inhibitory inputs, while the putative glutamatergic neurons receive more excitatory inputs in the reconstructed than in the randomized brain.

As a comparison, we also plotted the distributions of the E-I index for the randomized fruit fly brain network (see Methods). We found that the distributions of weighted inputs for the reconstructed fruit fly brain were much wider than those of the randomized one, suggesting that the neural connections in the fruit fly brain are organized in a way that leads to numerous neurons with high or low E-I index. This trend was much more significant for the weighted than the unweighted inputs. Specifically, we found that the putative cholinergic neurons (Cha) in the reconstructed brain are characterized by a wider and roughly symmetric distribution of the E-I index. In other words, this neuron population had equally large percentages of neurons with very high or very low E-I indices. In contrast, the putative GABArgic inhibitory neurons (Gad) in the reconstructed brain were characterized by a wider but asymmetric distribution of the E-I index, which indicates that there were many more Gad neurons receiving strong inhibitory input in the fruit fly brain network than in a randomized brain network. Moreover, the putative glutamatergic neurons (VGlu) in the reconstructed brain were characterized by a trend opposite to that of the inhibitory neurons: the VGlu neurons tend to receive stronger excitatory input than the inhibitory ones. One may suspect that the wide E-I index distributions of the reconstructed brain may had been artifacts due to subsampling of the full brain network. To address this question, we hypothesized that the full brain network (estimated to possess 100,000 - 150,000 neurons) is random-network like and exhibits narrow E-I index distributions, which become significantly widened after subsampling. We tested this hypothesis by constructing a random network of 130,000 neurons with the percentage of each neuron type and their connectivity (in percentage) following those in the reconstructed brain. Next, we randomly selected ~20,000 glutamatergic, cholinergic, and GABAergic neurons and calculated their E-I indices. We found that the subsampled random network exhibits much narrower E-I index distributions than those of the reconstructed brain (S2 Fig). Therefore, the hypothesis of subsampling artifacts was rejected.

The wide E-I index distributions of the reconstructed brain indicate that it is potentially unstable due to mutually suppressed inhibitory neurons and mutually facilitated excitatory neurons. Next, we investigated the actual stability of the fruit fly brain network by computer simulation.

### Dynamical properties of the fruit fly brain model

We performed the neural network simulations for the fruit fly brain model. At this early stage of whole-brain model development, we focused on establishing a stable resting state (see Methods) and on investigating its dynamical properties. The stability of the network is determined by the network structure and the overall strength of the excitatory and inhibitory connections. While the network structure was derived and determined by the connectomic data, the strength of the excitatory and inhibitory connections can be tuned by adjusting the variable *B* in Eq. 6. We defined the I/E factor as the ratio between *B*’s for the inhibitory and excitatory synapses. *B* was a fixed value (= 2.2) for all excitatory synapses, and therefore the I/E factor was determined by setting *B* for inhibitory synapses. For example, *B* is equal to *22* for the inhibitory synapses if the I/E factor is 10. We found that if we set the I/E factor to be one, the network was extremely unstable; the mean firing rate of the whole network quickly arose to nearly 100 Hz within 1 s. Next, we tuned the I/E factor and examined whether the brain network can be stabilized with a larger I/E factor (Figure 6). We varied the factor in the range between 0.1 and 100, and found that although the average firing rate of the whole network decreased dramatically with the increase of the I/E factor (Figure 6A), the network was still unstable. The instability was indicated by seizure-like firing activity, or hyperactivity, which is defined as a rapid surge of the mean firing rate of the whole brain to more than 1.0 Hz. Increasing I/E factor moderately prolonged the onset of the seizure-like events, but did not completely eliminate them (Figure 6B). We found that once the seizure-like activity occurred, it never stopped (Figures 6C and 6D). We further checked the distribution of the firing rate of individual neurons and found that for the case of I/E factor = 1, there were numerous neurons exhibiting extremely high firing rates (Figure 6E). While for a large I/E factor (100, for example), although the number of high firing rate neurons decreased, the distribution still exhibited a long tail (Figure 6F). The inefficiency of the I/E factor in stabilizing the network may be contributed by the following two factors: 1) Some of excitatory neurons have a highly positive E-I index, or less inhibitory input, making them less sensitive to strong inhibitory synapses. 2) The negative mean E-I index in the inhibitory neurons indicates strongly recurrent inhibition. Therefore, these neurons tend to inhibit themselves and limit the overall inhibitory output to the excitatory neurons.

**Figure 6.**
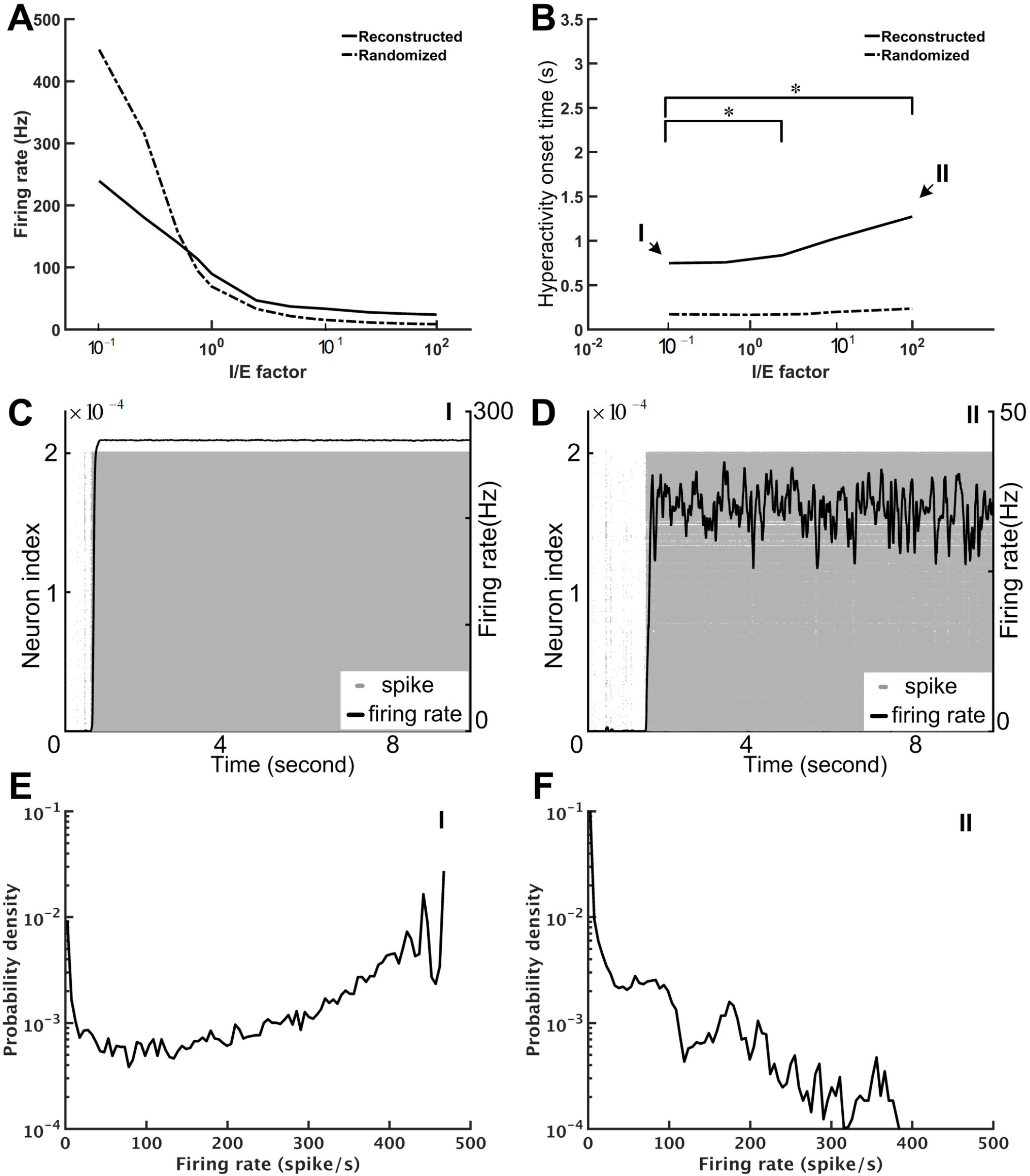
The seizure-like hyperactivity can be reduced but not completely eliminated by stronger weights for the inhibitory synapses, as represented by the I/E factor. (A) The mean firing rate as a function of the I/E factor for the reconstructed fruit fly brain and a randomized fruit fly brain. (B) The mean onset time of the seizure-like activity as a function of the I/E factor. Larger I/E factors significantly delay the onset time for the reconstructed fruit fly brain, but not for the randomized fruit fly brain. Furthermore, the reconstructed fruit fly brain is more stable than the randomized brain as indicated by the larger onset time for all I/E factors. Asterisks indicate the statistical significance (Student-t test, p<0.05) in the change of mean onset time between different I/E factor conditions for the reconstructed brain. (C) and (D) The spike rastergrams (grey dots) and the firing rates (black curves) of the reconstructed fruit fly brain at the low (0.1, point I in B) and high (100, point II in B) I/E factors, respectively. (E) and (F) The distributions of single neuron firing rates of the reconstructed fruit fly brain with the same I/E ratio as in (C) and (D), respectively.

Our simulations indicated that a strong inhibitory system, as characterized by a large I/E factor, is unable to stabilize the brain network. Therefore, we needed another neural mechanism that can efficiently “cool down” the network when the overall activity was high. To this end, we tested the short-term depression (STD), which is commonly observed in many species, including the *Drosophila*(Kazama and Wilson 2008). We implemented STD in every synapse of the fruit fly brain network and set the I/E factor equal to 10. We noted that the precise value of the I/E factor is not crucial. Setting the value above 5 would lead to the same network dynamics, qualitatively. We represented the degree of stability by the prevalence of the hyperactivity, as defined by its total duration in a 10-s simulation period, for different STD strengths, which is indicated by the recovery time constant (*τ_D_*) of STD. We found that STD effectively stabilized the reconstructed brain network and the prevalence dropped to 50% or lower when *τ_D_* was larger than 125 ms (Figure 7A). Moreover, while the seizure-like activity ran continuously in the brain network without STD (Figure 7B; Video S1), these hyperactivity events generally did not last for more than a few seconds in the brain network with strong STD (Figures 7B–D, Videos S2 and S3). This is intriguing considering that STD was not able to stabilize the randomized fly brain with *τ_D_* up to 1,000 ms (Figure 7A). When the hyperactivity was suppressed by a strong STD (*τ_D_* = 600 ms) in a reconstructed fly brain, it exhibited more diverse firing activity, as characterized by intermittent low activity and bursts of spikes with various durations (Figure 7D).

STD effectively stabilized the brain activity in terms of the population (the whole brain) firing rate. Next, we examined the activity of individual neurons by plotting the distribution of their mean firing rates. We found that although both reconstructed and randomized brain networks were characterized by broad firing rate distributions and could be fitted by power-law functions with exponential cut-off (*y* = A*x^−α^e^−βx^* or, truncated power law), they exhibited distinct characteristics (Figure 8). The firing rates distribution of the randomized brain network could also be fitted by an exponential function (Figure 8A) with small χ^2^ errors (~10^-3^), comparable to those in the fitting with a truncated power law (χ^2^~10^-3^). Moreover, the fitting with the truncated power law gave rise to an extremely small power-law exponent (*α*~0.012 − 0.058), indicating the insignificance of the power-law component in the distributions. Fitting the distributions with a power-law function alone yielded larger χ^2^ errors (~10^-1^-10^-2^). Therefore, we concluded that the firing rate distributions of the randomized brain network were better described by exponential functions. In contrast, the firing rate distributions of the reconstructed brain were better described by power-law than by exponential functions. Fitting the distributions with an exponential function did not yield any meaningful result (χ^2^ > 5.1) while fitting with a truncated power law distribution led to much smaller χ^2^ errors (~10^-2^-10^-3^). Furthermore, the power-law component was much more significant (*α*~1.48 − 0.84) in the reconstructed than in the randomized brain network.

**Figure 7.**
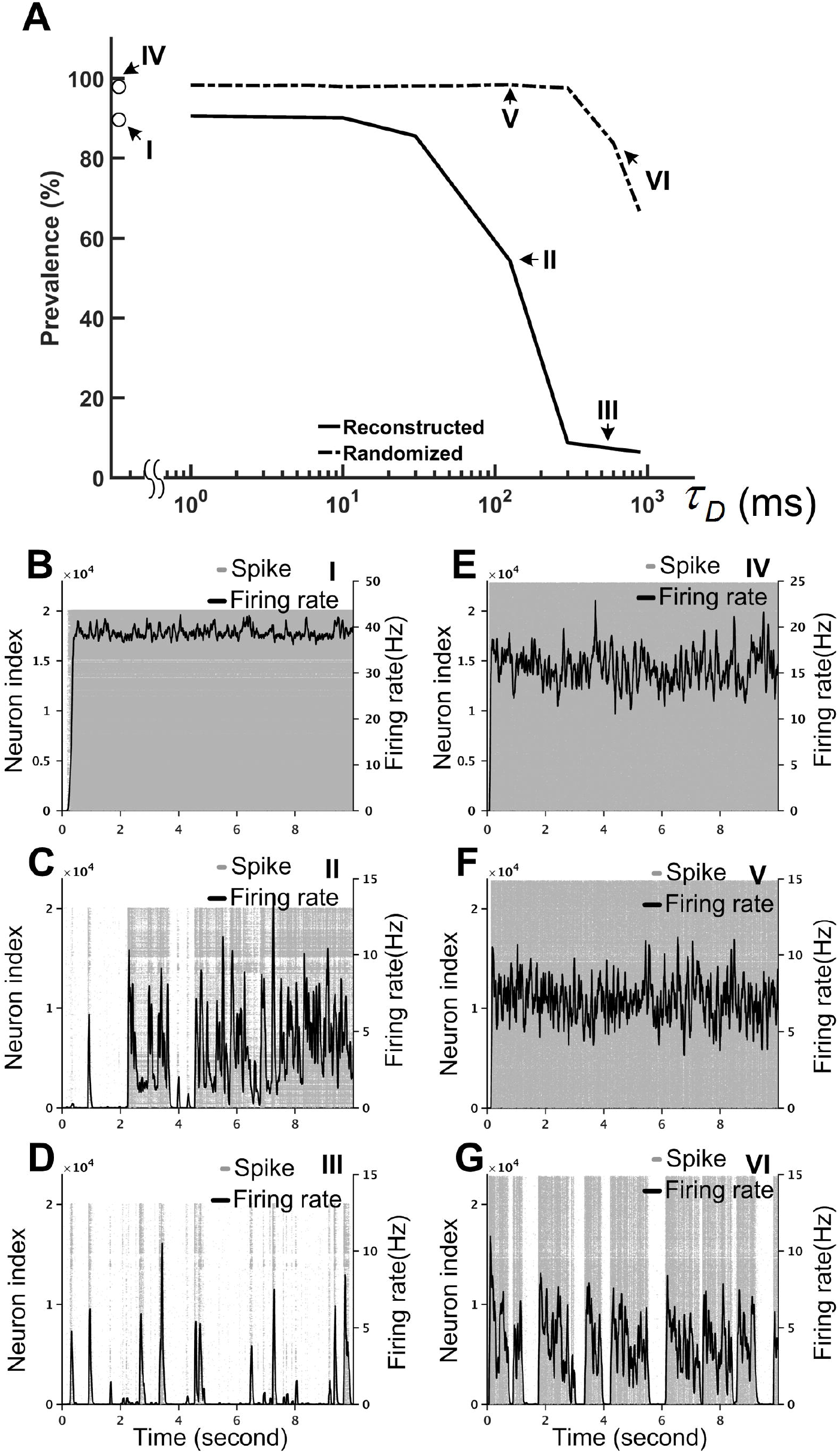
Short-term depression (STD) effectively stabilized the reconstructed fruit fly brain network by suppressing hyperactivity. The I/E factor is 10 in all panels. (A) The prevalence of hyperactivity, as defined by its total duration in a 10-s simulation period, as a function of the time constant of STD. A large time constant indicates stronger STD, which dramatically reduces the prevalence of hyperactivity for the reconstructed brain, but not for the randomized brain. (B) - (D) The spike rastergrams (grey dots) and the averaged firing rates (black curves) of the reconstructed fly brain without STD, with *τ_D_*=125 ms, and with *τ_D_*=600 ms, respectively. (E) - (G) Same as in (B) - (D), but for the randomized brain. The activity displayed in panels (B) - (G) corresponds to the data points labeled by the roman numerals I–VI in the panel (A), respectively.

So far, we have examined the mean neuronal activity at the population level (Figures 8 and 9) and at the single neuron level (Figure 8). In addition to the mean activity, the fluctuation of neuronal activity also exhibited distinct differences between the reconstructed brain network and the randomized one. We calculated the Fano factor for each neuron (10 trials, each lasting for 10 s) in the reconstructed and randomized networks (Figure 9) and found that while the mean Fano Factor was comparable between the two networks, the former had a much wider distribution than the latter. The result indicates that the reconstructed brain had highly diverse neural dynamics, characterized by a large number (compared to the randomized network) of neurons that fired randomly or with some non-random patterns. Intriguingly, we discovered that some of the high Fano factor neurons exhibited brief and high frequency burst activity with relatively long quiescent duration. Since the neurons were modeled with the simple leaky integrate-and-fire (LIF) model, such patterned activities were the result of network interactions.

**Figure 8.**
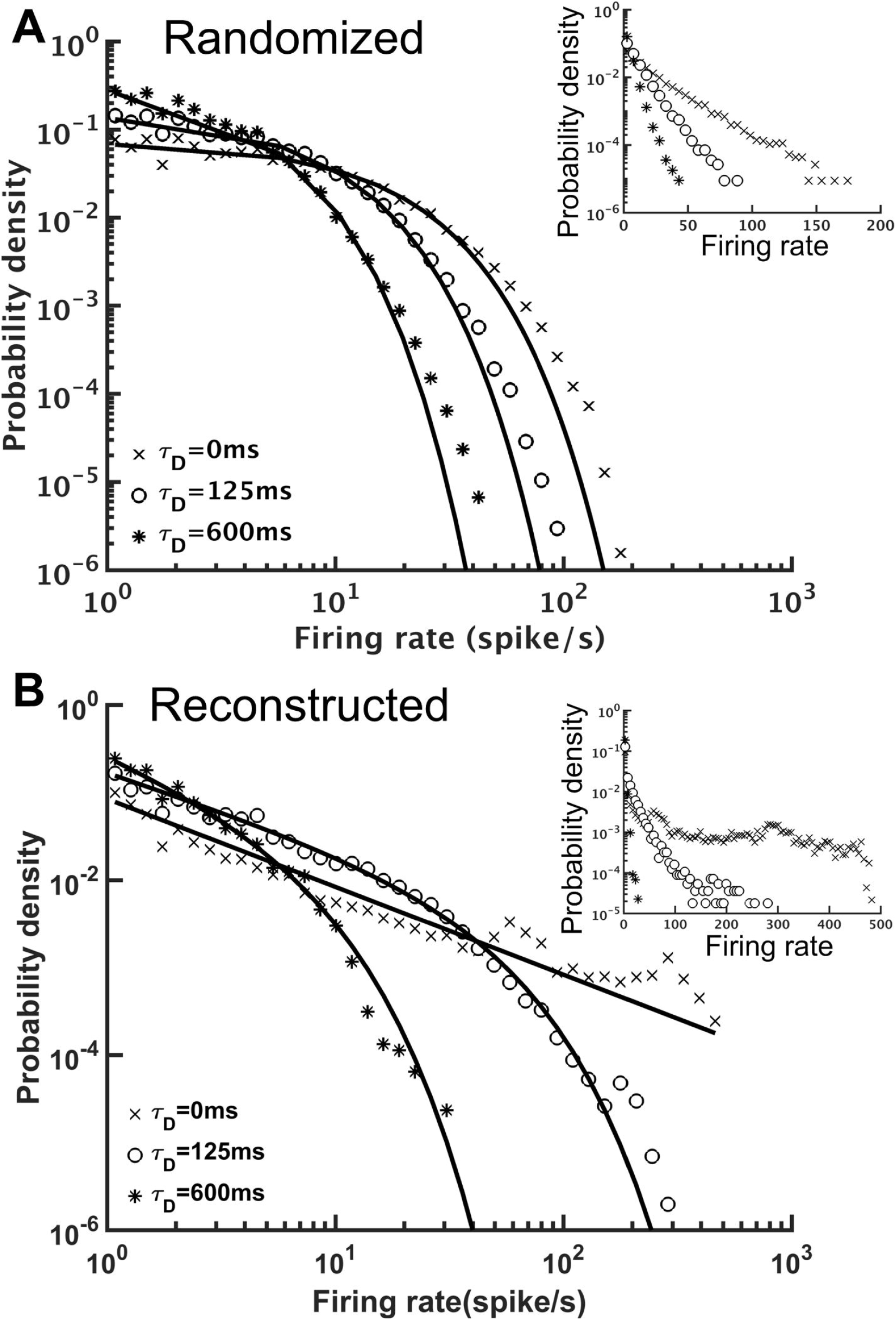
The distributions of single neuron mean firing rate with different short-term depression (STD) conditions. (A) The distributions for the randomized brain network with or without STD (*τ_D_* = 125 or 600 ms) in double-log plot. Inset, same data but in a semi-log plot. The solid lines indicate exponential fits to the distributions and the characteristic time constant decreases with. (B) Same as in (A), but for the reconstructed brain network. The distributions had strong power-law components and could be better fitted with a truncated power-law function (solid lines). Inset, same data but in a semi-log plot. The result indicated distinct dynamics between the randomized and reconstructed brain networks.

**Figure 9.**
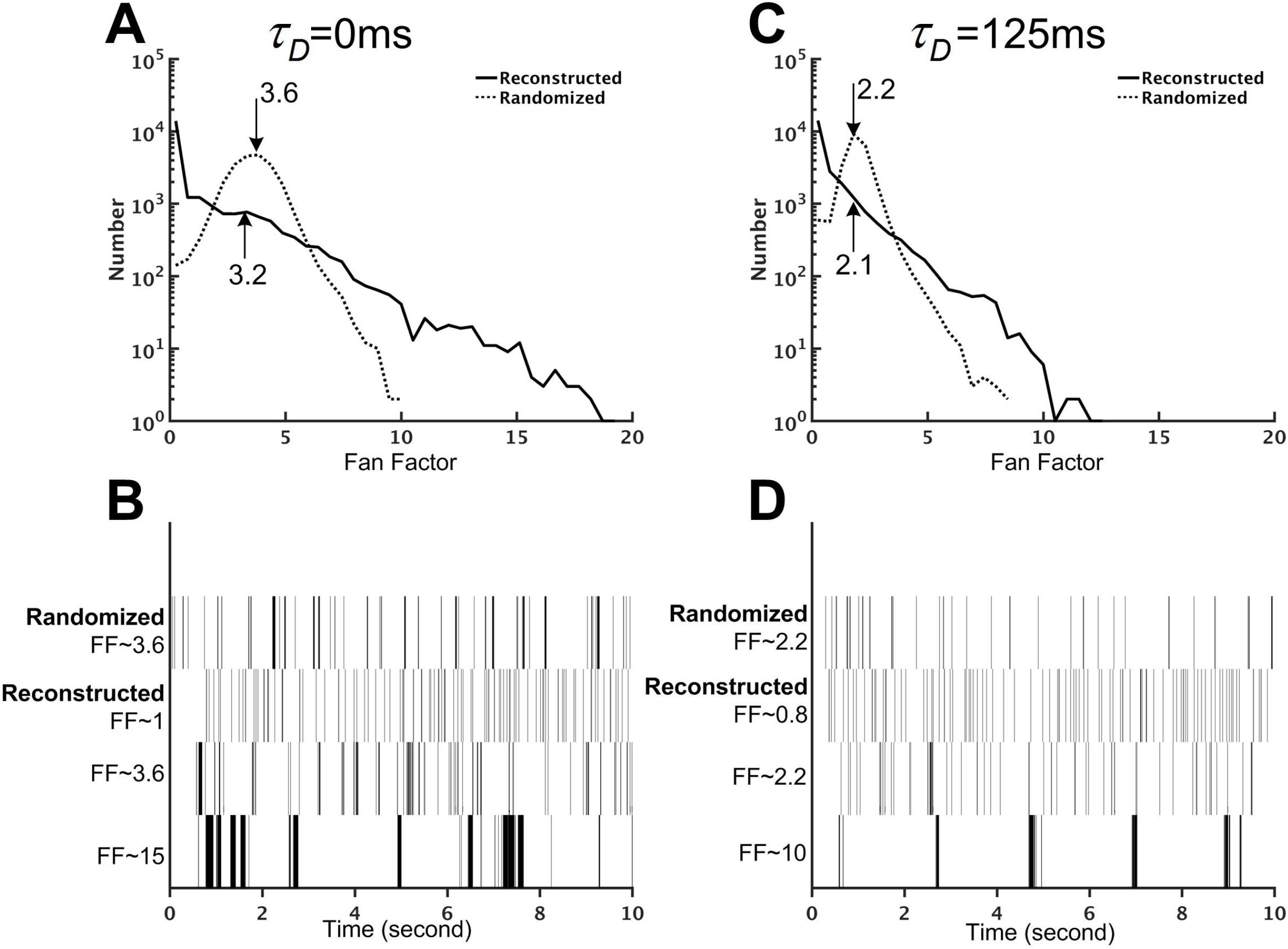
The reconstructed brain network exhibits firing patterns that are more diverse than those of the randomized brain network. (A) The distribution of the Fano factors for the reconstructed (solid line) and randomized (dotted line) brain networks without short-term depression (STD) (*τ_D_* = 0 ms). The arrows and the numbers indicate the mean of each distribution. Although both networks have comparable mean Fano factors, the reconstructed brain is characterized by a much wider distribution than the randomized one. (B) Sample spike trains with different Fano factors from the randomized and the reconstructed brain networks without STD (*τ_D_* = 0 ms). Neurons with high Fano factors in the reconstructed brain are characterized by clusters of bursting activity, while neurons with low Fano factors have a more evenly distributed spike activity. (C) Same as in (A) but with strong STD (*τ_D_* = 125 ms). (D) Same as in (B) but with strong STD (*τ_D_* = 125 ms).

### Simulator benchmark

We tested the performance of the Flysim simulator on a PC equipped with an Intel CPU at 3.6 GHz (E3-1270v5) with 64 Gigabytes of RAM. The reconstructed brain network (20,098 neurons and 1,044,020 synapses) required only 35 Mbytes of RAM and its simulation could be carried out in Flysim with four parallel threads at the speed of 1/35 of the real time. Next, we compared the Flysim simulator with NEST, a popular neural network simulator, using a simple 2-population random network. In the network, an excitatory population, E, of 16,000 neurons formed a recurrent circuit with an inhibitory population, I, of 4,000 neurons. The in-degree was set to 50 for each neuron. NEST provides a variety of neuron and synapse models. However, because the available combination of the neuron and synapse models do not exactly match those used in the Flysim simulator, we tested NEST with two sets of combinations, with one requiring more and the other requiring less computational power than our simulator. We first tested the HT model(Hill and Tononi 2005) in NEST because this model offers a synaptic dynamic that is comparable to that used in Flysim. However, the HT model is endowed with soma dynamics that are more complex than ours. Next, we also tested the LIF model, which is endowed with a simpler synapse model (iaf_psc_exp_multisynapse). The LIF model is comparable to ours but the synapse model is much simpler than that used in Flysim. Our result indicated that Flysim required less memory and ran faster than NEST in all conditions we tested (S3 Fig). We did not test NEST with the fruit fly brain model because it contains descriptions of parameters for more than 20,000 unique neurons and 1,000,000 unique synapses. The python interface used in NEST could not handle such a large size parameter input.

## Discussion

In the present study, we constructed the first brain-wide computational model based on the cellular-level connectome of the *Drosophila*. This model is the first of its kind for any species, except for *C. elegans*(Szigeti et al. 2014; Izquierdo and Beer 2016; Palyanov et al. 2011), which, however, is not considered to possess a brain. The proposed fly brain model, although still in its early stage of development, already exhibits several intriguing dynamical properties when compared to a randomized brain network. First, the E-I index was more widely distributed in the reconstructed brain network than in the randomized one, suggesting large populations of neurons receiving strongly excitatory or inhibitory inputs in the reconstructed brain. Second, despite the diversity in the E-I index, the reconstructed brain network was more stable, as measured by the prevalence of hyperactivity, than the randomized brain network. Third, although being more stable, the reconstructed fruit fly brain was characterized by diverse firing patterns: some neurons exhibited clusters of bursting activity while others fired more evenly.

The ultimate goal of our study is to develop a single-neuron level computational model of the fruit fly brain that can reproduce the detailed neuronal activity and behavior of fruit flies and that can be used to elucidate the computational principles of a fruit fly brain. Achieving such a goal requires a long-term effort together with highly detailed connectome and physiological data that are not yet available. Nevertheless, the purpose to present our early effort toward this goal in this paper is (1) to demonstrate, at the whole brain level, the unique dynamical features of a brain model reconstructed from the single-cell level connectome, and (2) by actually building one, to identify the technology and methodology that are required to improve the accuracy of the model, and (3) to draw attention to the issue regarding what exactly an “accurate brain model” means. We discuss these points as follows.

We demonstrated that both reconstructed and randomized networks are unstable at any level of the I/E factor without STD, and the reconstructed brain only becomes significantly more stable and diverse than the randomized one when STD is implemented. Therefore, the critical factor that leads to the stability of the reconstructed brain should be a certain interaction between the network structure and STD. It will be interesting to investigate which aspects of the network structure, globally or locally, may play roles in the STD-induced stability and study whether such structure characteristics exist in the brains of all species. Our study also delivered an important message: it is crucial to use a network structure that resembles a real brain. Using random networks, which are very popular among many theoretical studies of neural network dynamics, may not reveal the phenomena that actually occur in the brain.

The proposed fly brain model can be improved in several aspects:

1. Neuron type identification. Currently neuron types, including glutamatergic, GABAergic, and cholinergic, are recognized by the three GAL4 drivers, vGlut, GAD, and Cha, respectively. This driver-type mapping is known to be less than 100% accurate. Moreover, some neurons were found to release more than one types of neurotransmitters. Therefore, improved genetic tools are required in order to obtain more accurate cell type categorization (Diao et al. 2015).
2. Receptor type identification. Being a glutamatergic or GABAergic neuron does not automatically imply that the downstream neurons receive excitatory or inhibitory input, respectively. For example, glutamate-gated chloride channels have been observed in fruit flies. Since this type of channels cause an opposite effect to the AMPA and NMDA channels, it is important to conduct a systematic and high-resolution mapping of the expression of the synaptic receptors in the fruit fly brain so that the model can be updated accordingly.
3. Models of modulatory synapses. We currently only model four types of synaptic receptors: AMPA, NMDA, GABA_A_, and acetylcholine, which are fast-acting excitatory or inhibitory receptors. Therefore, the proposed fly brain circuits can only be used for model brain dynamics in short (sub-second) time scales. We will implement other slow-acting modulatory receptors, such as dopamine and serotonin, which will expect to endow the brain model with long-term and more complex behavior.
4. Polarity identification. The polarity of each neuron arbor is identified by the SPIN method. Although being highly efficient and reasonably accurate, the method still has room to improve. In particular, due to the small sizes and irregular morphology of local neurons, their polarity is more difficult to be correctly identified. Moreover, some local neurons have been shown to exhibit co-localized presynaptic and postsynaptic terminals (Chou et al. 2010). Improved image segmentation and tracing algorithms will provide more detailed morphological features for SPIN and will greatly improve its accuracy.
5. Image alignment and warping. Due to the potential deformation of the brain during the image acquisition process, when warping each neuron image into the standard brain space, it inevitably introduces errors that cause inaccuracy in the connection prediction. This issue will be largely improved by the *in situ* imaging method that will be adopted for the next generation of the FlyCircuit database. In addition, GRASP and related technology (Feinberg et al. 2008; Macpherson et al. 2015) can be used to verify the synapses and their activity in the selected circuits.
6. Single neuron model. Currently we use the single compartmental leaky integrate-and-fire model, and the membrane area is simply considered to be proportional to a neuron’s total branch length. As the information about the thickness of each branch will soon be available in the database, we will be able to more accurately calculate the area of the membrane and thus derive better estimates for related parameters. Adopting a multi-compartmental model will also help to improve the accuracy of the simulations (Günay et al. 2015). Moreover, some neurons in the visual system conduct signals by graded potentials or by mixed graded and action potentials (Mu et al. 2012; Baden et al. 2013). Although in the current study we only investigated the resting state activity of the model brain without visual stimulus, it is important to identify those non-spiking neurons in our sample and choose models that correctly represent their response properties in the future study which involves visual responses of the brain.

Finally, it is natural to ask how accurate the brain model is and how it can be verified. We would like to stress that, the term “accuracy” itself is not well-defined because of inter-individual differences. In the FlyCircuit database, each neuron image was taken from a different brain. Therefore, the reconstructed brain based on the database can be treated as an “average brain” sampled from a large number of individuals. In this sense, it is not meaningful to verify our fly brain model against a connectome reconstructed from a single brain. However, we argue that it is more meaningful to verify our brain model at the functional level; although each fruit fly may have slightly different brain circuits, they all perform the same basic functions. Although the connectome reconstructed based on electron microscopy has the potential to accurately reflect the neural network of one individual, it is not clear whether a model built upon one individual brain has an advantage over that built upon an average brain from the perspective of computer modeling. Moreover, an important perspective came from the consideration of neurodegenerative diseases, such as Alzheimer’s disease, which is characterized by significant loss of neurons and synapses. Unless in the advanced stages, patients with Alzheimer’s still maintain basic motor and cognitive functions, suggesting that these functions are robust against moderate alternation of neural circuits. Therefore, even though it is not possible to know whether the reconstructed fruit fly brain accurately reproduces the brain of any individual, as long as we continuously update the model with the availability of new data and improve the algorithms for estimating the model parameters, we presume that the reconstructed fruit fly brain will exhibit some basic brain functions in the near future.

Among all the brain functions, response to sensory input is the most suitable one for validating our brain model. In the next phase of model development, we will start with some of the most robust innate behaviors, such as the escape response, in which fruit flies jump directly away from a looming threat (von Reyn et al. 2017). A looming threat can be simulated by presenting a booming visual stimulus on the small field neurons in the unilateral medulla, while the initiation of the escape behavior can be represented by the activation of the giant fiber neurons (Tanouye and Wyman 1980). By then, the fruit fly brain model will provide an excellent platform for studying the neural circuit mechanisms of brain functions and behaviors.

## Acknowledgement

We thank National Center for High-performance Computing for providing computational resources. The work was supported by the Ministry of Science and Technology grants 105-2633-B-007-001 and 105-2311-B-007-012-MY3, and by the Aim for the Top University Project of the Ministry of Education.

## Author contribution

YCH wrote the paper, performed simulations and analyses. CTW performed simulations and analyses, and constructed the network model. TSS performed neural polarity prediction. KWK prepared figures. YJL and ASC provided the original single neuron data. CCL designed the study and wrote the paper.

## Information Sharing Statement

Flysim is an open-source neural network simulator released under the GNU General Public License (GPL v2+) and is available for download at https://github.com/yc-h/flysim.git. The neuronal data, the derived model parameters and the network connectivity are available at http://flycircuit.tw upon the publication of this paper.

---

**S1 Fig. The input (ordinate) and output (abscissa) contact points of each neuron generally follow a linear relationship in a double-logarithmic plot** The solid line represents the linear regression of the data: log(y) = 0.48 * log(x) + 2.6.

**S2 Fig. E-I index distributions for a subsampled network in comparison to the reconstructed and randomized fruit fly brain networks** To test whether the broad E-I index distributions of the reconstructed brain network are artifacts due to subsampling from the full brain network, we constructed a full-size (130,000 neurons) random network (see text) and subsampled it by randomly selecting 22,835 neurons from the full network. The subsampled network exhibits much narrower E-I index distributions than those of the reconstructed brain network.

**S3 Fig. Benchmark tests indicated the superior efficiency and performance of the Flysim simulator compared to a similar simulator** (A) To perform the comparison, we constructed a simple recurrent network with two populations. The network has 20,000 neurons and each one receives input from 50 randomly chosen neurons. Population E consists of 16,000 excitatory neurons while population I contains 4,000 GABAergic neurons. (B) We recorded that memory usage and the performance, as measured by the amount of CPU time required to simulate one second of biological time, for the Flysim simulator and NEST v2.12.0. For NEST, we tested the HT and leaky integrate-and-fire (LIF) models. In all conditions, the Flysim simulator consumed less memory and performed faster than NEST.

**S1 Table. Summary of membrane properties of neurons in *Drosophila* and the source of the data** These values are used to determine the standard membrane properties in the model (see text).

**S1 Video. Activity of the reconstructed fruit fly brain without short-term depression**

**S2 Video. Activity of the reconstructed fruit fly brain with moderate short-term depression (*τ_D_* = 125 ms)**

**S3 Video. Activity of the reconstructed fruit fly brain with strong short-term depression (*τ_D_* = 600 ms)**

## Reference

Abbott, L. F., J. A. Varela, Kamal Sen, and S. B. Nelson. 1997. “Synaptic Depression and Cortical Gain Control.” Science 275 (5297): 221–24. https://doi.org/10.1126/science.275.5297.221.

Baden, Tom, Thomas Euler, Matti Weckström, and Leon Lagnado. 2013. “Spikes and Ribbon Synapses in Early Vision.” Trends in Neurosciences 36 (8): 480–88. https://doi.org/10.1016/j.tins.2013.04.006.

Berg, Rune W, Aidas Alaburda, and Jørn Hounsgaard. 2007. “Balanced Inhibition and Excitation Drive Spike Activity in Spinal Half-Centers.” Science 315 (5810): 390–93. https://doi.org/10.1126/science.1134960.

Burns, Randal, Joshua T. Vogelstein, and Alexander S. Szalay. 2014. “From Cosmos to Connectomes: The Evolution of Data-Intensive Science.” Neuron 83 (6): 1249–52. https://doi.org/10.1016/j.neuron.2014.08.045.

Chance, Frances S, L F Abbott, and Alex D Reyes. 2002. “Gain Modulation from Background Synaptic Input.” Neuron 35 (4): 773–82. https://doi.org/10.1016/S0896-6273(02)00820-6.

Chang, Po-Yen, Ta-Shun Su, Chi-Tin Shih, and Chung-Chuan Lo. 2017. “The Topographical Mapping in Drosophila Central Complex Network and Its Signal Routing.” Frontiers in Neuroinformatics 11. https://doi.org/10.3389/fninf.2017.00026.

Chaudhuri, Rishidev, and Ila Fiete. 2016. “Computational Principles of Memory.” Nature Neuroscience 19 (3): 394–403. https://doi.org/10.1038/nn.4237.

Chiang, Ann-Shyn, Chih-Yung Lin, Chao-Chun Chuang, Hsiu-Ming Chang, Chang-Huain Hsieh, Chang-Wei Yeh, Chi-Tin Shih, et al. 2011. “Three-Dimensional Reconstruction of Brain-Wide Wiring Networks in Drosophila at Single-Cell Resolution.” Current Biology 21 (1): 1–11. https://doi.org/10.1016/j.cub.2010.11.056.

Chou, Ya-Hui, Maria L Spletter, Emre Yaksi, Jonathan C S Leong, Rachel I Wilson, and Liqun Luo. 2010. “Diversity and Wiring Variability of Olfactory Local Interneurons in the Drosophila Antennal Lobe.” Nature Neuroscience 13 (4): 439–49. https://doi.org/10.1038/nn.2489.

Churchland, Anne K., and L. F. Abbott. 2016. “Conceptual and Technical Advances Define a Key Moment for Theoretical Neuroscience.” Nature Neuroscience 19 (3): 348–49. https://doi.org/10.1038/nn.4255.

Cox, S. M., and P. C. Matthews. 2002. “Exponential Time Differencing for Stiff Systems.” Journal of Computational Physics 176 (2): 430–55. https://doi.org/10.1006/jcph.2002.6995.

Cuntz, Hermann, Friedrich Forstner, Alexander Borst, and Michael Häusser. 2010. “One Rule to Grow Them All: A General Theory of Neuronal Branching and Its Practical Application.” PLoS Comput Biol 6 (8): e1000877. https://doi.org/10.1371/journal.pcbi.1000877.

Denève, Sophie, and Christian K. Machens. 2016. “Efficient Codes and Balanced Networks.” Nature Neuroscience 19 (3): 375–82. https://doi.org/10.1038/nn.4243.

Diao, Fengqiu, Holly Ironfield, Haojiang Luan, Feici Diao, William C. Shropshire, John Ewer, Elizabeth Marr, Christopher J. Potter, Matthias Landgraf, and Benjamin H. White. 2015. “Plug-and-Play Genetic Access to Drosophila Cell Types Using Exchangeable Exon Cassettes.” Cell Reports 10 (8): 1410–21. https://doi.org/10.1016/j.celrep.2015.01.059.

Douglass, John K., and Nicholas J. Strausfeld. 2003. “Anatomical Organization of Retinotopic Motion-Sensitive Pathways in the Optic Lobes of Flies.” Microscopy Research and Technique 62 (2): 132–150. https://doi.org/10.1002/jemt.10367.

Eliasmith, Chris, Terrence C. Stewart, Xuan Choo, Trevor Bekolay, Travis DeWolf, Yichuan Tang, and Daniel Rasmussen. 2012. “A Large-Scale Model of the Functioning Brain.” Science 338 (6111): 1202–5. https://doi.org/10.1126/science.1225266.

Fawcett, Tom. 2006. “An Introduction to ROC Analysis.” Pattern Recognition Letters, ROC Analysis in Pattern Recognition, 27 (8): 861–74. https://doi.org/10.1016/j.patrec.2005.10.010.

Feinberg, Evan H., Miri K. VanHoven, Andres Bendesky, George Wang, Richard D. Fetter, Kang Shen, and Cornelia I. Bargmann. 2008. “GFP Reconstitution Across Synaptic Partners (GRASP) Defines Cell Contacts and Synapses in Living Nervous Systems.” Neuron 57 (3): 353–63. https://doi.org/10.1016/j.neuron.2007.11.030.

“Frontiers | Neuroarch: A Graph-Based Platform for Constructing and Querying Models of the Fruit Fly Brain Architecture.” n.d. Accessed January 18, 2017. http://www.frontiersin.org/10.3389/conf.fninf.2014.18.00042/event_abstract.

Givon, Lev E., and Aurel A. Lazar. 2016. “Neurokernel: An Open Source Platform for Emulating the Fruit Fly Brain.” PLOS ONE 11 (1): e0146581. https://doi.org/10.1371/journal.pone.0146581.

Gouwens, Nathan W., and Rachel I. Wilson. 2009. “Signal Propagation in Drosophila Central Neurons.” The Journal of Neuroscience 29 (19): 6239–49. https://doi.org/10.1523/JNEUROSCI.0764-09.2009.

Günay, Cengiz, Fred H. Sieling, Logesh Dharmar, Wei-Hsiang Lin, Verena Wolfram, Richard Marley, Richard A. Baines, and Astrid A. Prinz. 2015. “Distal Spike Initiation Zone Location Estimation by Morphological Simulation of Ionic Current Filtering Demonstrated in a Novel Model of an Identified Drosophila Motoneuron.” PLOS Computational Biology 11 (5): e1004189. https://doi.org/10.1371/journal.pcbi.1004189.

Haider, B, A Duque, A R Hasenstaub, and D A McCormick. 2006. “Neocortical Network Activity in Vivo Is Generated through a Dynamic Balance of Excitation and Inhibition.” J Neurosci, 4535–45. https://doi.org/10.1523/JNEUROSCI.5297-05.2006.

Helmstaedter, Moritz. 2013. “Cellular-Resolution Connectomics: Challenges of Dense Neural Circuit Reconstruction.” Nature Methods 10 (6): 501–7. https://doi.org/10.1038/nmeth.2476.

Hempel, Chris M., Kenichi H. Hartman, X.-J. Wang, Gina G. Turrigiano, and Sacha B. Nelson. 2000. “Multiple Forms of Short-Term Plasticity at Excitatory Synapses in Rat Medial Prefrontal Cortex.” Journal of Neurophysiology 83 (5): 3031–41.

Hill, Sean, and Giulio Tononi. 2005. “Modeling Sleep and Wakefulness in the Thalamocortical System.” J Neurophysiol 93 (3): 1671–1698. https://doi.org/10.1152/jn.00915.2004.

Izhikevich, Eugene M., and Gerald M. Edelman. 2008. “Large-Scale Model of Mammalian Thalamocortical Systems.” Proceedings of the National Academy of Sciences 105 (9): 3593–98. https://doi.org/10.1073/pnas.0712231105.

Izquierdo, Eduardo J, and Randall D Beer. 2016. “The Whole Worm: Brain–Body–Environment Models of C. Elegans.” Current Opinion in Neurobiology, Systems neuroscience, 40 (October): 23–30. https://doi.org/10.1016/j.conb.2016.06.005.

Kazama, Hokto, and Rachel I. Wilson. 2008. “Homeostatic Matching and Nonlinear Amplification at Identified Central Synapses.” Neuron 58 (3): 401–13. https://doi.org/10.1016/j.neuron.2008.02.030.

Lasko, Thomas A., Jui G. Bhagwat, Kelly H. Zou, and Lucila Ohno-Machado. 2005. “The Use of Receiver Operating Characteristic Curves in Biomedical Informatics.” Journal of Biomedical Informatics, Clinical Machine Learning, 38 (5): 404–15. https://doi.org/10.1016/j.jbi.2005.02.008.

Lee, Yi-Hsuan, Yen-Nan Lin, Chao-Chun Chuang, and Chung-Chuan Lo. n.d. “SPIN: A Method of Skeleton-Based Polarity Identification for Neurons.” Neuroinformatics, 1–21. https://doi.org/10.1007/s12021-014-9225-6.

Lin, Chih-Yung, Chao-Chun Chuang, Tzu-En Hua, Chun-Chao Chen, Barry J. Dickson, Ralph J. Greenspan, and Ann-Shyn Chiang. 2013. “A Comprehensive Wiring Diagram of the Protocerebral Bridge for Visual Information Processing in the Drosophila Brain.” Cell Reports 3 (5): 1739–53. https://doi.org/10.1016/j.celrep.2013.04.022.

Lo, Chung-Chuan, and Ann-Shyn Chiang. 2016. “Toward Whole-Body Connectomics.” Journal of Neuroscience 36 (45): 11375–83. https://doi.org/10.1523/JNEUROSCI.2930-16.2016.

Lo, Chung-Chuan, Cheng-Te Wang, and Xiao-Jing Wang. 2015. “Speed-Accuracy Tradeoff by a Control Signal with Balanced Excitation and Inhibition.” Journal of Neurophysiology, May, jn.00845.2013. https://doi.org/10.1152/jn.00845.2013.

Macpherson, Lindsey J., Emanuela E. Zaharieva, Patrick J. Kearney, Michael H. Alpert, Tzu-Yang Lin, Zeynep Turan, Chi-Hon Lee, and Marco Gallio. 2015. “Dynamic Labelling of Neural Connections in Multiple Colours by Trans-Synaptic Fluorescence Complementation.” Nature Communications 6 (December): 10024. https://doi.org/10.1038/ncomms10024.

Mariño, Jorge, James Schummers, David C Lyon, Lars Schwabe, Oliver Beck, Peter Wiesing, Klaus Obermayer, and Mriganka Sur. 2005. “Invariant Computations in Local Cortical Networks with Balanced Excitation and Inhibition.” Nature Neuroscience 8 (2): 194–201. https://doi.org/10.1038/nn1391.

Markram, Henry. 2006. “The Blue Brain Project.” Nature Reviews Neuroscience 7 (2): 153–60. https://doi.org/10.1038/nrn1848.

Milham, Michael Peter. 2012. “Open Neuroscience Solutions for the Connectome-Wide Association Era.” Neuron 73 (2): 214–18. https://doi.org/10.1016/j.neuron.2011.11.004.

Morante, Javier, and Claude Desplan. 2008. “The Color-Vision Circuit in the Medulla of Drosophila.” Current Biology 18 (8): 553–65. https://doi.org/10.1016/j.cub.2008.02.075.

Morgan, Joshua L., and Jeff W. Lichtman. 2013. “Why Not Connectomics?” Nature Methods 10 (6): 494–500. https://doi.org/10.1038/nmeth.2480.

Mu, Laiyong, Kei Ito, Jonathan P. Bacon, and Nicholas J. Strausfeld. 2012. “Optic Glomeruli and Their Inputs in Drosophila Share an Organizational Ground Pattern with the Antennal Lobes.” Journal of Neuroscience 32 (18): 6061–71. https://doi.org/10.1523/JNEUROSCI.0221-12.2012.

Nagel, Katherine I., Elizabeth J. Hong, and Rachel I. Wilson. 2015. “Synaptic and Circuit Mechanisms Promoting Broadband Transmission of Olfactory Stimulus Dynamics.” Nature Neuroscience 18 (1): 56–65. https://doi.org/10.1038/nn.3895.

Osumi-Sutherland, David, Simon Reeve, Christopher J. Mungall, Fabian Neuhaus, Alan Ruttenberg, Gregory S. X. E. Jefferis, and J. Douglas Armstrong. 2012. “A Strategy for Building Neuroanatomy Ontologies.” Bioinformatics 28 (9): 1262–69. https://doi.org/10.1093/bioinformatics/bts113.

Palyanov, Andrey, Sergey Khayrulin, Stephen D. Larson, and Alexander Dibert. 2011. “Towards a Virtual C. Elegans: A Framework for Simulation and Visualization of the Neuromuscular System in a 3D Physical Environment.” In Silico Biology 11 (3–4): 137–47. https://doi.org/10.3233/ISB-2012-0445.

Parekh, Ruchi, and Giorgio A. Ascoli. 2013. “Neuronal Morphology Goes Digital: A Research Hub for Cellular and System Neuroscience.” Neuron 77 (6): 1017–38. https://doi.org/10.1016/j.neuron.2013.03.008.

Peng, Hanchuan, Michael Hawrylycz, Jane Roskams, Sean Hill, Nelson Spruston, Erik Meijering, and Giorgio A. Ascoli. 2015. “BigNeuron: Large-Scale 3D Neuron Reconstruction from Optical Microscopy Images.” Neuron 87 (2): 252–56. https://doi.org/10.1016/j.neuron.2015.06.036.

Peters, Alan, and Bertram R. Payne. 1993. “Numerical Relationships between Geniculocortical Afferents and Pyramidal Cell Modules in Cat Primary Visual Cortex.” Cerebral Cortex 3 (1): 69–78. https://doi.org/10.1093/cercor/3.1.69.

Reyn, Catherine R. von, Aljoscha Nern, W. Ryan Williamson, Patrick Breads, Ming Wu, Shigehiro Namiki, and Gwyneth M. Card. 2017. “Feature Integration Drives Probabilistic Behavior in the Drosophila Escape Response.” Neuron 94 (6): 1190–1204.e6. https://doi.org/10.1016/j.neuron.2017.05.036.

Root, Cory M., Julia L. Semmelhack, Allan M. Wong, Jorge Flores, and Jing W. Wang. 2007. “Propagation of Olfactory Information in Drosophila.” Proceedings of the National Academy of Sciences 104 (28): 11826–31. https://doi.org/10.1073/pnas.0704523104.

Shinomiya, Kazunori, Keiji Matsuda, Takao Oishi, Hideo Otsuna, and Kei Ito. 2011. “Flybrain Neuron Database: A Comprehensive Database System of the Drosophila Brain Neurons.” The Journal of Comparative Neurology 519 (5): 807–33. https://doi.org/10.1002/cne.22540.

Shu, Yousheng, Andrea Hasenstaub, and David A. McCormick. 2003. “Turning on and off Recurrent Balanced Cortical Activity.” Nature 423 (6937): 288–93. https://doi.org/10.1038/nature01616.

Sporns, Olaf. 2013. “Making Sense of Brain Network Data.” Nature Methods 10 (6): 491–93. https://doi.org/10.1038/nmeth.2485.

Su, Ta-Shun, Wan-Ju Lee, Yu-Chi Huang, Cheng-Te Wang, and Chung-Chuan Lo. 2017. “Coupled Symmetric and Asymmetric Circuits Underlying Spatial Orientation in Fruit Flies.” Nature Communications 8 (1): 139. https://doi.org/10.1038/s41467-017-00191-6.

Szigeti, Balázs, Padraig Gleeson, Michael Vella, Sergey Khayrulin, Andrey Palyanov, Jim Hokanson, Michael Currie, Matteo Cantarelli, Giovanni Idili, and Stephen Larson. 2014. “OpenWorm: An Open-Science Approach to Modeling Caenorhabditis Elegans.” Frontiers in Computational Neuroscience 8: 137. https://doi.org/10.3389/fncom.2014.00137.

Takemura, Shin-ya, Arjun Bharioke, Zhiyuan Lu, Aljoscha Nern, Shiv Vitaladevuni, Patricia K. Rivlin, William T. Katz, et al. 2013. “A Visual Motion Detection Circuit Suggested by Drosophila Connectomics.” Nature 500 (7461): 175–81. https://doi.org/10.1038/nature12450.

Tanaka, Nobuaki K., Keita Endo, and Kei Ito. 2012. “Organization of Antennal Lobe-Associated Neurons in Adult Drosophila Melanogaster Brain.” Journal of Comparative Neurology 520 (18): 4067–4130. https://doi.org/10.1002/cne.23142.

Tanouye, M A, and R J Wyman. 1980. “Motor Outputs of Giant Nerve Fiber in Drosophila.” Journal of Neurophysiology 44 (2): 405–21. https://doi.org/10.1152/jn.1980.44.2.405.

“The Ziggurat Method for Generating Random Variables | Marsaglia | Journal of Statistical Software.” n.d. Accessed March 24, 2017. https://www.jstatsoft.org/article/view/v005i08.

Ukani, Nikul H., Chung-Heng Yeh, Adam Tomkins, Yiyin Zhou, Dorian Florescu, Carlos Luna Ortiz, Yu-Chi Huang, et al. 2016. “The Fruit Fly Brain Observatory: From Structure to Function.” BioRxiv, December, 092288. https://doi.org/10.1101/092288.

Varela, Juan A., Kamal Sen, Jay Gibson, Joshua Fost, L. F. Abbott, and Sacha B. Nelson. 1997. “A Quantitative Description of Short-Term Plasticity at Excitatory Synapses in Layer 2/3 of Rat Primary Visual Cortex.” Journal of Neuroscience 17 (20): 7926–40.

Vogels, Tim P, and L F Abbott. 2009. “Gating Multiple Signals through Detailed Balance of Excitation and Inhibition in Spiking Networks.” Nat Neurosci 12 (4): 483–91. https://doi.org/10.1038/nn.2276.

Wang, Cheng-Te, Chung-Ting Lee, Xiao-Jing Wang, and Chung-Chuan Lo. 2013. “Top-Down Modulation on Perceptual Decision with Balanced Inhibition through Feedforward and Feedback Inhibitory Neurons.” PLoS ONE 8 (4): e62379. https://doi.org/10.1371/journal.pone.0062379.

Wang, X.-J. 1999. “Fast Burst Firing and Short-Term Synaptic Plasticity: A Model of Neocortical Chattering Neurons.” Neuroscience 89 (2): 347–62. https://doi.org/10.1016/S0306-4522(98)00315-7.

Webb, Barbara, and Antoine Wystrach. 2016. “Neural Mechanisms of Insect Navigation.” Current Opinion in Insect Science 15 (June): 27–39. https://doi.org/10.1016/j.cois.2016.02.011.

Weir, Peter T., Bettina Schnell, and Michael H. Dickinson. 2014. “Central Complex Neurons Exhibit Behaviorally Gated Responses to Visual Motion in Drosophila.” Journal of Neurophysiology 111 (1): 62–71. https://doi.org/10.1152/jn.00593.2013.

Wessnitzer, Jan, and Barbara Webb. 2006. “Multimodal Sensory Integration in Insects—towards Insect Brain Control Architectures.” Bioinspiration & Biomimetics 1 (3): 63. https://doi.org/10.1088/1748-3182/1/3/001.

Wilson, Rachel I., and Gilles Laurent. 2005. “Role of GABAergic Inhibition in Shaping Odor-Evoked Spatiotemporal Patterns in the Drosophila Antennal Lobe.” J. Neurosci. 25 (40): 9069–79.

Wolff, Tanya, Nirmala A. Iyer, and Gerald M. Rubin. 2015. “Neuroarchitecture and Neuroanatomy of the Drosophila Central Complex: A GAL4-Based Dissection of Protocerebral Bridge Neurons and Circuits.” Journal of Comparative Neurology 523 (7): 997–1037. https://doi.org/10.1002/cne.23705.

Zheng, Zhihao, J. Scott Lauritzen, Eric Perlman, Camenzind G. Robinson, Matthew Nichols, Daniel Milkie, Omar Torrens, et al. 2018. “A Complete Electron Microscopy Volume of the Brain of Adult Drosophila Melanogaster.” Cell 174 (3): 730–743.e22. https://doi.org/10.1016/j.cell.2018.06.019.

Zhu, Yan. 2013. “The Drosophila Visual System.” Cell Adhesion & Migration 7 (4): 333–44. https://doi.org/10.4161/cam.25521.

